# Locus-specific spatial and transcriptional dynamics direct T cell lineage choice in the thymus

**DOI:** 10.64898/2026.10.26.761611

**Authors:** Ilinca Patrascan, James WD King, Noura Maziak, Rebecca M Crossley, Samuel Johnson, Arianna Ceccarelli, Ian McFarlane, Ya Guo, Mohammad M Karimi, Amanda G Fisher, Ruth E Baker, Juan M Vaquerizas, Matthias Merkenschlager

## Abstract

Decades of investigation have made the choice between CD4 and CD8 T cell lineages in the thymus a key paradigm for mammalian cell commitment and differentiation1-29, yet how transcriptional diversification is coordinated with 3D genome organisation remains largely unexplored. Here, using single-molecule transcript quantification and region capture Pico-C (RCPC), we resolve distinct transcriptional and spatial chromatin dynamics at the lineage-defining loci Cd4 and Cd8. We find early Cd4/Cd8 transcriptional divergence driven by MHC-dependent bias in Cd8a transcript copy number, which unfolds gradually across both CCR7− and CCR7+ differentiation stages. Unexpectedly, Cd4 and Cd8 loci show fundamentally different spatial contact dynamics: transiently reduced mesoscale (10-100kb) contacts at Cd8 provide a permissive chromatin environment throughout the transcriptional diversification process. By contrast, Cd4 undergoes switch-like contact changes limited to the microscale (1kb) late in differentiation, associated with lineage stabilization by ThPOK and/or RUNX3. These results place distinct, locus-specific chromatin environments at the centre of transcriptional diversification during T cell lineage choice.

## Introduction

The choice between the CD4 and the CD8 T cell lineage in the thymus is a classic paradigm of developmental cell fate commitment and the dynamic regulation of the lineage-defining loci *Cd4* and *Cd8* is central to this process^1–29^. T cell progenitors rearrange and express clone-specific T cell receptors (clonotypic TCRs) that recognise antigens presented by major histocompatibility complex (MHC) molecules, a property known as MHC restriction^1,2^. CD4 and CD8 are coreceptors for TCR engagement with MHC class II or MHC class I, respectively. Co-expression of *Cd4* and *Cd8* in bipotent thymic progenitors is resolved into mutually exclusive expression to align co-receptor expression with TCR specificity and CD4 T_helper_ or CD8 T_cytotoxic_ gene expression programmes^1,2,3,5,6,19^ (Supplementary Fig. 1a). Although the signalling logic of CD4/CD8 lineage choice has been studied extensively^5,11,17–20,24–29^, it remains unclear when transcriptional divergence at *Cd4* and *Cd8* loci first emerges and how this divergence is coordinated with changes in chromatin architecture during lineage commitment.

A central unresolved issue is whether lineage choice is established through early branching or through a shared intermediate phase followed by later divergence. One-step models posit that MHC class I- and class II-signalled progenitors diverge soon after TCR engagement^7,9,10^ (Supplementary Fig. 1b), whereas sequential models propose an initial common differentiation path marked by termination of *Cd8* transcription and a later bifurcation associated with re-expression of *Cd8* in MHC class I-restricted cells^3,4,6,17,18,20^ (Supplementary Fig. 1c). Recent single-cell transcriptomic analyses have refined this framework^20–23^, but questions remain about the timing, progression, and mechanistic basis of transcriptional asymmetry at the lineage-defining loci.

In addition, little is known about how 3D chromatin organisation at the *Cd4* and *Cd8* loci changes during the developmental transition from CD4 CD8 double positive (DP) progenitors towards CD4 or CD8 single positive (SP) progeny, or whether these loci are regulated through shared or distinct structural principles. This is especially important because lineage choice requires not only changes in gene expression, but the stabilisation of alternative regulatory states at two opposing loci within the same differentiating cell. Defining how transcriptional diversification is related to locus-specific chromatin organisation is therefore central to understanding how lineage choice is executed.

Here, we combine single-molecule RNA fluorescence in situ hybridisation to quantify nascent and mature *Cd4* and *Cd8* transcripts in defined thymocyte intermediates with low-input, ultra-high-resolution RCPC to map 3D contacts across the *Cd4* and *Cd8* loci *in vivo* (Supplementary Fig. 1d). We show that transcriptional divergence begins earlier and progresses more gradually than predicted by strictly sequential models, and that the two loci adopt distinct chromatin-organisational strategies during this process. Together, these results place locus-specific 3D chromatin environments at the centre of transcriptional diversification during T cell lineage choice.

## Results

### Tracking CD4/CD8 lineage choice and differentiation by smRNA-FISH

Prior to CD4/CD8 lineage choice, MHC-naive CD69^-^DP progenitors (CD69^-^DP) homogeneously express both *Cd4* and *Cd8a* (Fig. 1a, b left). Differentiation intermediates (CD69^+^ DP and CD4^+^8^lo^) show pronounced heterogeneity (Fig. 1a, b center), which resolves after CD4/CD8 lineage choice (Fig. 1a, b right). Transcriptional heterogeneity of differentiation intermediates is observed across multiple systems^30,31^. Sequential models predict such heterogeneity during CD4/CD8 lineage choice as CD69^+^ DP differentiation intermediates are thought to contain at least two distinct subsets of cells: i) early differentiation intermediates signaled by MHC class I or II prior to the CD4^+^8^lo^ stage and ii) late MHC class I-signaled differentiation intermediates that re-express CD8 subsequent to auditioning for the CD4 lineage at the CD4^+^8^lo^ stage and re-enter the DP population *en route* to becoming CD8 SP cells^3,4,6,17,18,20^ (Supplementary Fig. 1e). Sequential models also predict heterogeneity within the CD4^+^8^lo^ subset of differentiation intermediates, which has been proposed to arise as differentiation intermediates initially terminate CD8 transcription, even when destined for the CD8 lineage^17^. The chemokine receptors CCR4 and CCR7 are expressed sequentially in differentiation intermediates, CCR4 early and CCR7 later during CD4/CD8 lineage choice^32,33^. CCR7 is thought to demarcate a turning point where MHC class I-signaled differentiation intermediates undergo reversal of coreceptor expression following a failed attempt to adopt a CD4 fate^3,4,18^ (Supplementary Fig. 1e). However, it remains unclear exactly how and when *Cd4*/*Cd8* transcriptional diversification of differentiation intermediates occurs. We therefore set out to resolve the timing and trajectory of *Cd4/Cd8* transcriptional divergence by quantitative approaches at single-cell resolution across early and late differentiation intermediates.

**Figure 1.**
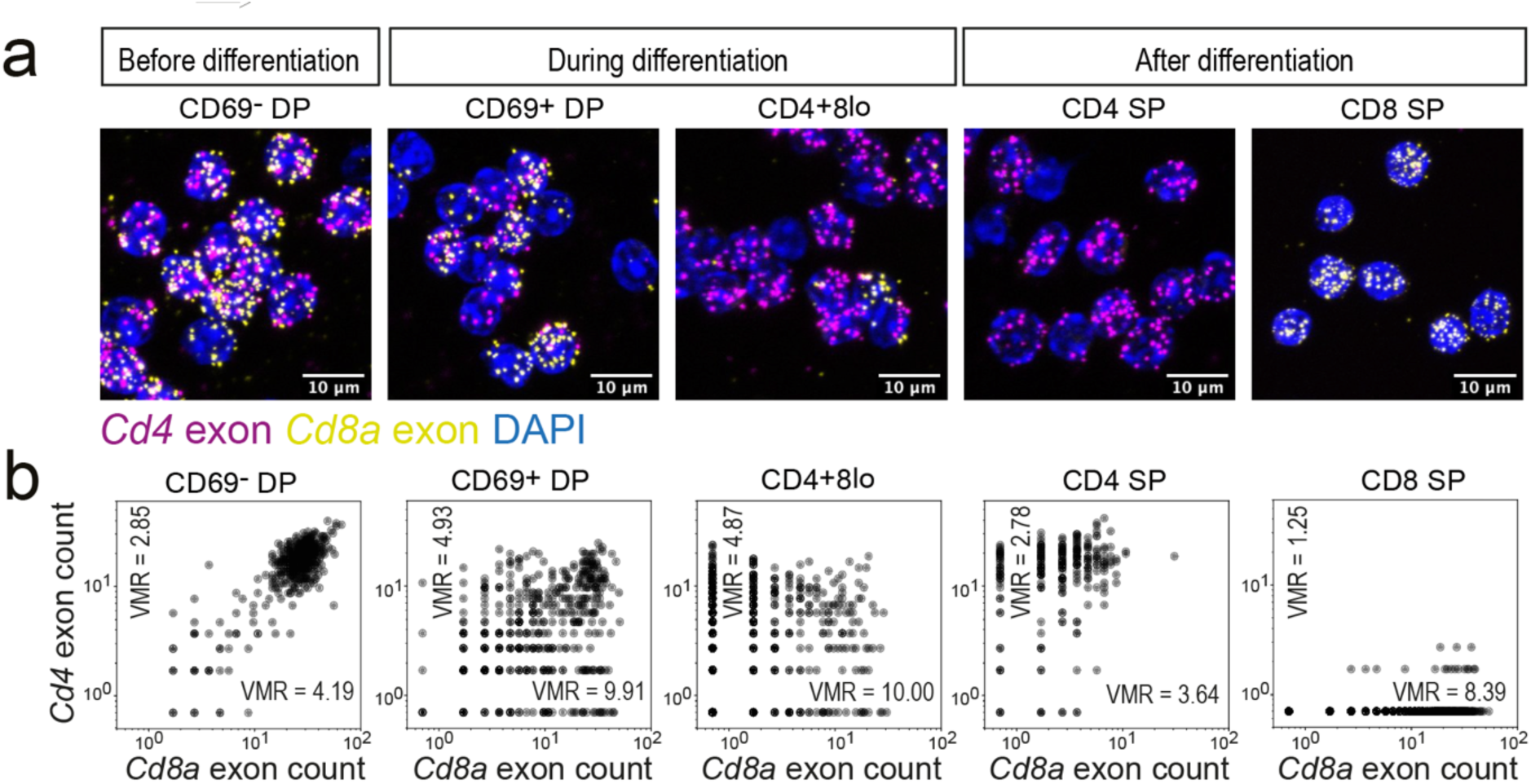
Tracking CD4/CD8 lineage choice and differentiation by single molecule RNA-FISH. a, Homogeneous expression of *Cd4* and *Cd8a* mRNA in bipotential progenitors (CD69^-^ DP) gives way to cell-to-cell heterogeneity in differentiation intermediates (CD69^+^ DP and CD4^+^8^lo^) and resolves after lineage choice (CD4 and CD8 SP). See Extended Data Fig. 1a for sorting strategy. b, *Cd4* and *Cd8a* mRNA copy number counts were determined by smRNA-FISH for the cell populations shown in a). VMR: variance to mean ratios, also known as index of dispersion (VMR = variance/mean) are shown for the X and Y axes. Representative of 2-4 independent experiments.

### CD8-fated differentiation intermediates initially downregulate and later restore *Cd8a* expression at the transcriptional level

To directly relate transcript copy number to CD4/CD8 lineage choice, it is important to know the ultimate fate of differentiation intermediates. A genetic approach to achieve this is to use Aβ^b-/-^(MHC class II-deficient) or *B2m*^-/-^(MHC class I-deficient) mice, where differentiation intermediates are signaled selectively either through MHC class I (class I-signaled differentiation intermediates) or through MHC class II (class II-signaled differentiation intermediates) - also referred to as CD8- or CD4-fated cells^6,20^. We used cell surface staining and flow cytometry to isolate class II- or class I-signaled early (CCR4^+^ CCR7^-^) and late (CCR7^+^) CD4^+^8^lo^ differentiation intermediates (Extended Data 1b). We then determined *Cd8a* intron and exon counts, *Cd4* intron counts and transcriptional burst counts for thousands of cells across multiple replicates by smRNA-FISH (Fig. 2a, Extended Data Fig. 2a-d for probe positions, probe specificity and the definition of transcriptional bursts). Pre-differentiation CD69^-^DP and post-differentiation CD4 SP and CD8 SP cells provided a framework for these analyses (Fig 2a; Supplementary Fig. 1d). Intronic transcript counts for the housekeeping gene *Actb* provided a reference for overall transcriptional activity during CD4/CD8 lineage choice and differentiation (Extended Data Fig. 2d).

**Figure 2.**
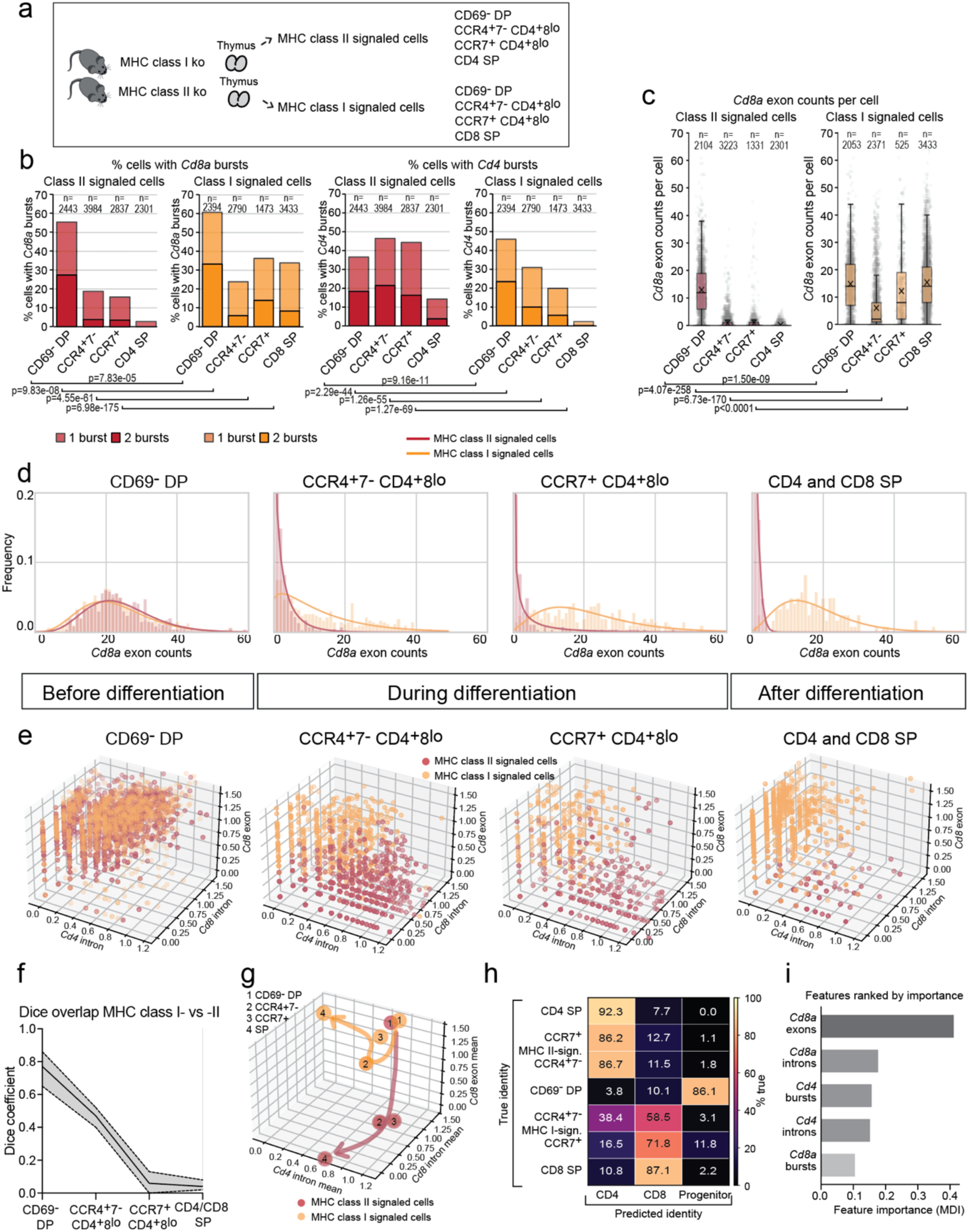
Early transcriptional diversification of CD4-versus CD8-fated differentiation intermediates. a, Experimental outline. The indicated populations were sorted from MHC class I- or MHC class II-deficient mice and subjected to smRNA-FISH. For some experiments, CD69^-^ DP progenitor cells were isolated from thymi of MHC class I/II double-deficient mice. See Extended Data Fig. 1b for sorting strategy. b, Percentage of cells within the indicated subsets that show transcriptional bursts for *Cd8a* (left) and *Cd4* (right). Stats: Chi square test. c, *Cd8a* exon counts for the indicated cell populations. The full data set, including *Cd8a* intron counts, *Cd4* intron counts and mean counts per replicate is shown in Extended Data Fig. 3a-c. n = the number of cells scored for each probe and cell population. X represents the mean in box plots throughout this manuscript. Stats: Chi square test. d, smRNA-FISH *Cd8a* exon counts (bars) and fitted two-species negative binomial models (lines) for the indicated cell populations. e, 3D plots of log *Cd8a* intron and exon and log *Cd4* intron counts. Note that transcriptional diversification is gradual, and visible prior to the acquisition of CCR7 f, Quantification of *Cd4/Cd8* divergence between MHC class I- and MHC class II-signaled differentiation intermediates using the Dice overlap or Dice Similarity Coefficient (using 90% kernel density estimates of the data shown in panel e. 50 bins and 200 bootstraps. g, 3D pathway schematic using log *Cd8a* intron and exon and log *Cd4* intron counts as a reference. h, Random forest classification of thymocytes using *Cd8a* exon and intron counts, *Cd4* intron counts, and *Cd8a* and *Cd4* transcriptional burst counts. Note that the classifier confidently predicts the fate of differentiation intermediates prior to the acquisition of CCR7. i) Relative feature importance of smRNA-FISH measurements in the random forest classifier. *Cd8a* mRNA copy number is most important for predicting lineage fate of differentiation intermediates.

We observed downregulation of *Cd8a* in differentiation intermediates signaled by either MHC class II or MHC class I (Fig. 2b, c, Extended Data Fig. 3a-c), even though differentiation intermediates signaled by MHC class I are ultimately destined for the CD8 lineage. *Cd8a* was subsequently re-expressed in MHC class I-signaled differentiation intermediates (Fig. 2b, c, Extended Data 3a-c).

Differentiation intermediates that express the high affinity class I-restricted OT-I TCR are known to form a clear CD4^+^8^lo^ population, whereas differentiation intermediates expressing other TCRs, such as the lower affinity F5 TCR, do not form a prominent CD4^+^8^lo^ population^18^ (Extended Data Fig. 4a). To address whether *Cd8a* downregulation would occur in MHC class I-signaled differentiation intermediates that do not transit through a CD4^+^8^lo^ stage, we isolated F5 differentiation intermediates. To accommodate the lack of CD4^+^8^lo^ cells, we adopted a previously validated classification based on CD69 and CCR7 (Ref.^18^; Extended Data Fig. 4a) and isolated OT-I differentiation intermediates using the same approach for comparison. smRNA-FISH with *Cd8a* intron (Extended Data Fig. 4b, c) and exon probes (Extended Data 4d) showed that both F5 and OT-I first downregulate and then re-induce *Cd8a* at the transcriptional level, even though F5 differentiation intermediates do not form a prominent CD4^+^8^lo^ population at the protein level.

Taken together, these data are broadly in line with the predictions of sequential (CD4-first) models^3,4,6^, with two important provisos. First, we extend sequential models by providing direct evidence for reversal of coreceptor expression. Second, we revise strictly sequential models by demonstrating important quantitative differences between MHC class I- and class II-signaled differentiation intermediates as described below.

### *Cd8a* is more strongly downregulated in MHC class II-signaled than in MHC class I-signaled differentiation intermediates

According to sequential models, both class I- and class II-signaled differentiation intermediates terminate *Cd8* transcription in response to TCR signals^17^. Our smRNA-FISH data support the view that MHC class II-signaled differentiation intermediates terminate *Cd8a* transcription. Importantly, however, MHC class I-signaled differentiation intermediates do not terminate but merely downregulate *Cd8* transcript levels. This observation is supported by qRT-PCR data on sorted thymocyte populations (Extended Data Fig. 3d). In our published scRNA-seq data obtained by full-length Smart-seq2 cDNA sequencing^22^, *Cd8a/b1* expression in MHC class I-signaled CD4^+^8^lo^ differentiation intermediates is several-fold higher than in MHC class I/II-signaled CD4^+^8^lo^ differentiation intermediates (*Cd8a* mean WT = 0.32, class II ko = 1.64; Extended Data Fig. 3e). Quantitative transcript analysis therefore revises the view of strictly sequential (CD4 first) models of CD4/CD8 lineage choice by demonstrating that the degree of *Cd8a* downregulation shows pronounced lineage-specific differences throughout (Fig. 2).

### Early transcriptional diversification of CD4-versus CD8-fated differentiation intermediates

The quantitative differences in the extent of *Cd8a* downregulation by TCR/co-receptor signaling through MHC class I versus MHC class II described here inform our view of CD4/CD8 lineage choice, as they show that the nature of TCR engagement imposes an early bias in *Cd8a* transcript copy numbers. Consequently, *Cd4/Cd8* transcriptional divergence between MHC class I- and II-signaled differentiation intermediates initiates early. smRNA-FISH shows that, following this initial bias, *Cd4/Cd8* transcriptional divergence evolves gradually and spans both CCR7^-^ and CCR7^+^ CD4^+^8^lo^ stages of CD4/CD8 lineage choice. Fig. 2e visualises the resulting changes across developmental stages. We quantified transcriptional divergence based on the Dice coefficient, which measures the overlap of *Cd8a* intron and exon and *Cd4* intron space between MHC class I- and II-signaled differentiation intermediates (see Methods). This demonstrated significant separation of MHC class I- and II-signaled differentiation intermediates from the CCR4^+^7^-^ CD4^+^8^lo^ stage onwards (Fig. 2f), preceding the acquisition of CCR7. A schematic of differentiation trajectories is provided in Fig. 2g.

### Random forest classifiers distinguish between CD4- and CD8-fated cells based on *Cd8a* transcript copy number

Machine learning confidently predicts the fate of differentiation intermediates prior to the acquisition of CCR7: a random forest classifier based on *Cd4* and *Cd8a* intron and *Cd8a* exon data (80% training, 20% testing) identified MHC class II-signaled differentiation intermediates as CD4-fated with high and comparable accuracy at both CCR4^+^CCR7^-^ (86.7%) and CCR7^+^ CD4^+^8^lo^ stages (86.2%). By contrast, the accuracy of classifying MHC class I-signaled differentiation intermediates as CD8-fated increased progressively from CCR4^+^CCR7^-^ (58.5%) to CCR7^+^ CD4^+^8^lo^ (71.8%) to CD8 SP (87.1%; Fig. 2h). Therefore, consistent with visual inspection (Fig. 2e) and statistical analysis of the data (Fig. 2f), the machine learning results indicate robust transcriptional diversification of differentiation intermediates prior to the acquisition of CCR7 (Fig. 2h). *Cd8a* mRNA copy number was the key parameter for the distinction between CD4- and CD8-fated differentiation intermediates by the machine learning approach (Fig. 2i).

F5 and OT-I TCR transgenic differentiation intermediates differed with respect to the developmental stage at which they could clearly be assigned to the CD8 lineage: the random forest classifier confidently assigned CD8 fate to F5 differentiation intermediates from the CD69^+^ CCR7^-^stage 2 onwards (Extended Data Fig. 4e), but remained undecided about the fate of OT-I differentiation intermediates until they acquired CCR7 expression (Extended Data Fig. 4e).

Hence, quantitative transcript analysis reveals how the onset of *Cd4/Cd8* transcriptional diversification initiates with an early bias in *Cd8a* downregulation imposed by TCR/co-receptor signaling through MHC class I versus MHC class II, respectively. Importantly, *Cd4/Cd8* symmetry is not abruptly broken, but rather dissolves gradually, spanning both CCR7^-^ and CCR7^+^ CD4^+^8^lo^ stages of CD4/CD8 lineage choice, and this process involves non-linearities particularly of *Cd8* expression. These findings add to an extensive body of work indicating that developing T cells must flexibly explore the *Cd4/Cd8* transcriptional space in order to align MHC restriction with co-receptor expression^1–4,19^ and raise intriguing questions about the 3D chromatin environments that underpin CD4/CD8 lineage choice.

### RCPC enables exploration of 3D chromatin environments that underpin CD4/CD8 lineage choice and differentiation

Elucidating 3D chromatin contacts that underpin CD4/CD8 lineage choice and differentiation represents a considerable challenge, as CD4/CD8 differentiation intermediates are rare cells that can be isolated in limited numbers only. We therefore built on Pico-C, a recently described and highly efficient Micro-C approach^34^, to develop low-input, ultra-high-resolution RCPC (Fig. 3a).

**Figure 3.**
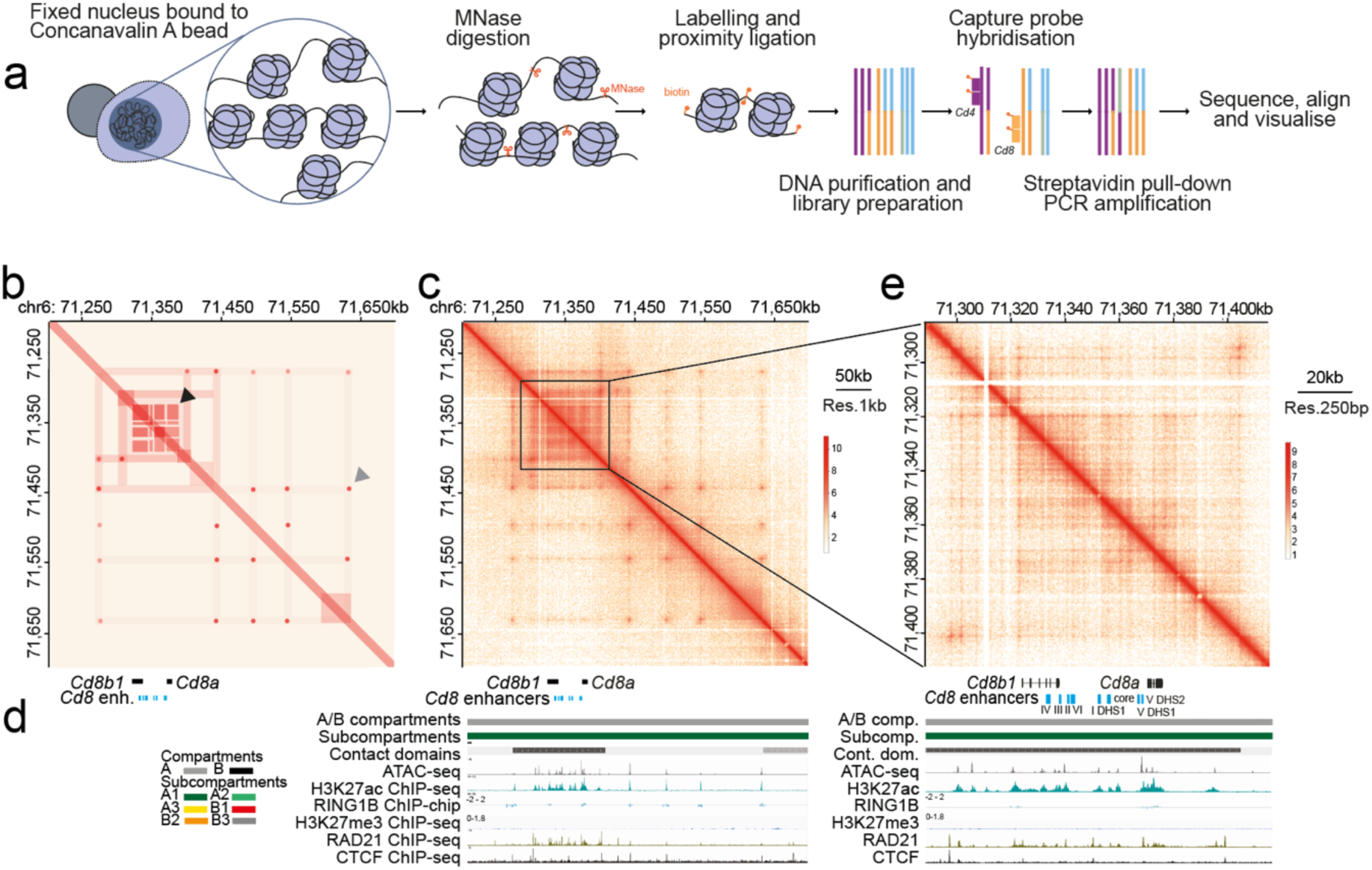
A characteristic checkerboard pattern marks *Cd8* gene-enhancer contacts. a, Outline of region capture Pico-C (RCPC) approach developed for this study b, Schematic representation of RCPC chromatin contacts in the *Cd8* capture region. Grey arrowhead: microcompartments. Black arrowhead: checkerboard-like contacts c, Merged RCPC data of the region depicted in b (count matrix 1kb resolution). Known enhancers^14^ are indicated. d, Chromatin features of the Cd8 capture region: Genes and enhancers (only genes within the *Cd8* core region are shown for clarity), A/B compartments and subcompartments^37^ - note that the entire *Cd8* region is classified as A1 subcompartment, Contact domains, ATAC-seq accessibility^38^, H3K27ac ChIP-seq (Ref.^37^), RING1B ChIP-chip (Ref.^39^), RAD21, CTCF and H3K27me3 ChIP-seq (Ref.^37^). e, View of the *Cd8* core region (count matrix 250bp resolution)

We prepared Pico-C libraries^34^ in two independent biological replicates each from bipotential CD69^-^ DP progenitors, CCR7^-^ and CCR7^+^ CD4^+^8^lo^ differentiation intermediates signaled by either MHC class I or MHC class II, and post-differentiation CD4 and CD8 SP populations (Fig. 2, Supplementary Fig. 1d). We designed oligonucleotides to capture ∼1Mb of sequence surrounding the *Cd4* and *Cd8* loci to sequence ∼30 million 150bp paired end reads, resulting in RCPC maps of a depth that would require roughly 8 billion reads when extrapolated to the whole genome (Extended Data Fig. 6a). Replicates across all biological conditions were of comparable depth and at least 85% of uniquely mapped reads were *in cis* (≥ 78% after deduplication, Extended Data Fig. 6a), indicating that the RCPC data are of high quality^34,35^.

### A characteristic checkerboard pattern marks contacts between *Cd8* genes and enhancers

The *Cd*8 core region was marked by a characteristic checkerboard-like contact pattern between gene bodies and regulatory elements^14^, whereas dot-like microcompartments familiar from previous capture Micro-C data^35^ featured prominently in the wider *Cd8* capture region (Fig. 3c-e). Checkerboard-like contacts have previously been described for target regions of Polycomb repressive (PRC) complexes^36^. However, the *Cd8* capture region is devoid of both the PRC1 component RING1B and the repressive histone mark H3K27me3 catalysed by PRC2 (Fig. 3d; the HoxA cluster on chromosome 6 serves as a positive control for RING1B and H3K27me3 in developing T cells, Extended Data Fig. 5). The *Cd8* capture region is assigned to the A compartment throughout, specifically the A1 subcompartment^37^. The observed checkerboard-like contacts correspond to the *Cd8* core region containing the *Cd8a* and *Cd8b1* coding genes and known *Cd8* enhancers^14^ (E8I – E8VI, Fig. 3b-e, Fig. 4a). The *Cd8* core region is marked by high levels of chromatin accessibility and the permissive histone mark H3K27ac (Fig. 3d), consistent with active transcription.

**Figure 4.**
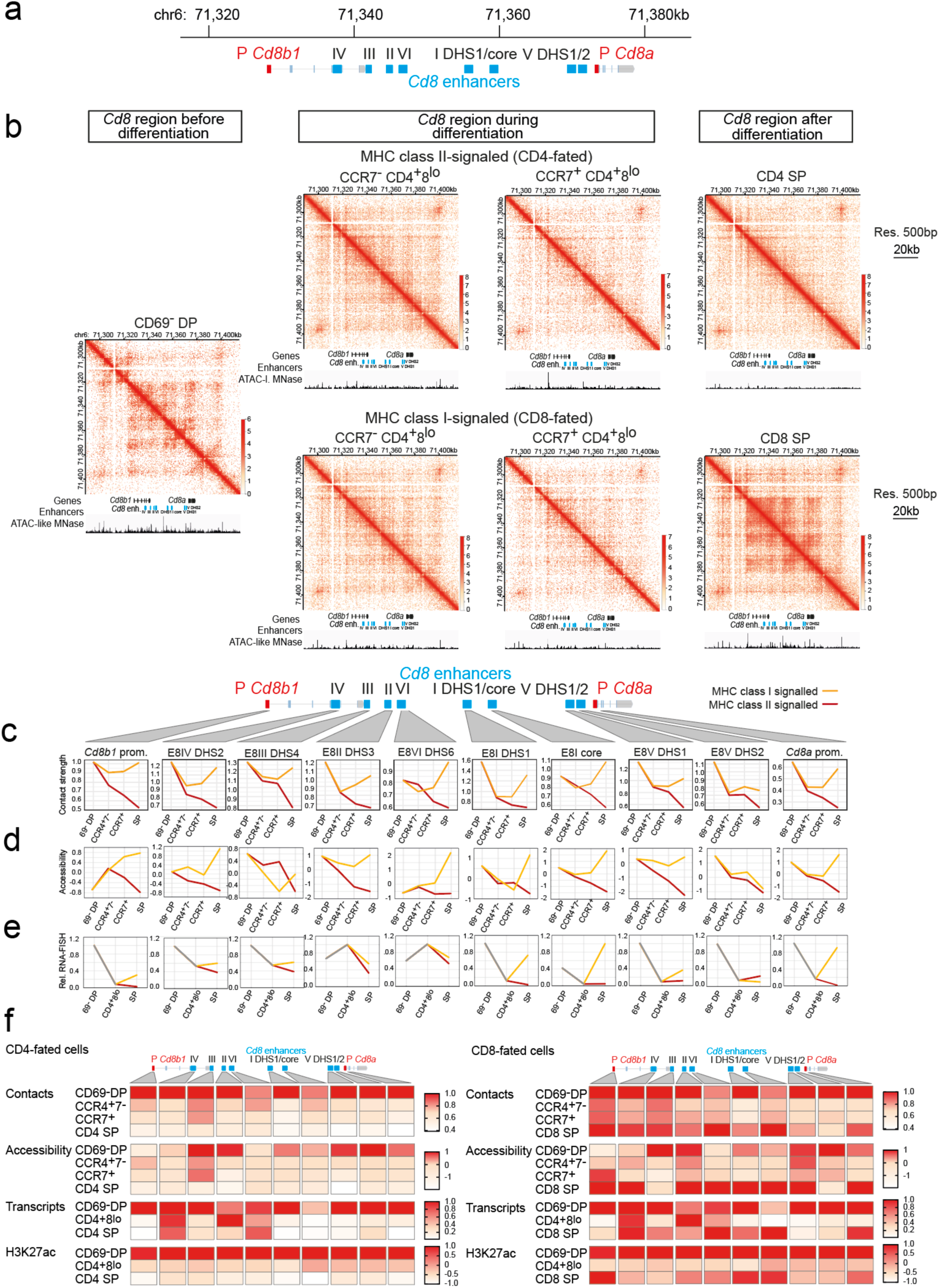
Transient weakening of *Cd8* gene-enhancer contacts in differentiation intermediates fated for either the CD4 or the CD8 lineage. a, Overview of the *Cd8* locus and its regulatory elements. P *Cd8a*: *Cd8a* promoter, P *Cd8b1*: promoter. *Cd8* enhancers are indicated in roman numerals I-VI (Ref.^14^) b, RCPC maps (count matrix 500bp resolution, natural log scale), 2 merged biological replicates. ATAC-like tracks below the contact heatmaps show accessibility based on analysis of <70bp MNase junction reads (See Methods and Extended Data Fig. 7c). Fig. 3 shows a larger view of the annotation tracks. c, Developmental regulation of total RCPC contacts made by each *Cd8* regulatory element with other *Cd8* regulatory elements. See Extended Data Fig. 7a, b for a fine-grained analysis how these contacts are distributed and reconfigured before, during and after CD4/CD8 lineage choice. d, Developmental regulation of ATAC-like MNase accessibility of *Cd8* regulatory elements e, Developmental regulation of genic and intergenic transcripts at the *Cd8* locus as quantified by smRNA-FISH f, Summary heatmap of contact and chromatin changes at the *Cd8* locus in cells fated for the CD4 (left) versus the CD8 lineage (right)

### Transient weakening of *Cd8* gene-enhancer contacts in differentiation intermediates

Visual inspection of RCPC data across developmental stages showed a striking pattern where *Cd8* enhancer-gene contacts are clearly defined in bipotential progenitors prior to CD4/CD8 lineage choice (Fig. 4a, b left) and become transiently attenuated in CCR7^-^ and CCR7^+^ differentiation intermediates (Fig. 4b, center) fated for both CD4 and CD8 lineages (Fig. 4b, center top for CD4-fated cells and center bottom for CD8-fated cells, respectively; Extended Data Fig. 6b schematically depicts the key changes in the chromatin contact maps). This attenuation of chromatin contacts in differentiation intermediates is selective for the ∼60kb core region containing *Cd8* genes and previously described enhancers^14^ because characteristic capture Micro-C microcompartments^35^ and other Micro-C features persisted in the wider capture region outside the *Cd8* core region throughout CD4/CD8 lineage choice and differentiation (Extended Data Fig. 6c). Residual contacts dissolve in CD4 SP cells (Fig. 4b, right top) but are restored in CD8 SP cells (Fig. 4b, right bottom).

Quantitative analysis of contact strength confirmed that reduced *Cd8* enhancer-gene contacts occurred irrespective of whether differentiation intermediates were signaled by MHC class I or MHC class II (Fig. 4b). Attenuation of *Cd8* enhancer-gene contacts was therefore independent of ultimate lineage fate. The developmental window of reduced *Cd8* enhancer-gene contacts spanned both the CCR7^-^ and CCR7^+^ CD4^+^8^lo^ stages (Fig. 4c). Residual *Cd8* enhancer-gene contacts dissipated in post-differentiation CD4 SP but were reinstated in CD8 SP (Fig. 4c). Fine-grained analysis showed substantial reconfiguration of contacts made by individual *Cd8* regulatory elements not just in differentiation intermediates, but also between CD69^-^ DP progenitor cells and mature CD8 SP cells (Extended Data Fig. 7a, b). Specifically, DHS1 and the core element of *Cd8* enhancer I gained interactions with the *Cd8a* and *Cd8b1* promoter regions in mature CD8 SP cells compared to progenitor cells prior to differentiation (Extended Data Fig. 7a, b). This is in line with selective activity of E8I in CD8 SP cells^14^. Hence, not only the overall strength, but also the distribution of contacts between *Cd8* regulatory elements are developmentally regulated. Pre-existing contacts in progenitor cells are weakened during CD4/CD8 lineage choice and reinstated in reconfigured form in mature CD8 lineage cells.

We utilised short (<70bp) junction reads as a proxy for MNase-based chromatin accessibility (See Methods and Extended Data Fig. 7c). Changes in the accessibility of *Cd8* regulatory elements across CD4/CD8 lineage choice followed a similar trend as contacts (Fig. 4d), with notable exceptions. For example, the accessibility of *Cd8* enhancer VI DHS-6 was low in CD69^-^ DP progenitors and increased steadily in MHC class I-but not in MHC class II-signaled differentiation intermediates (Fig. 4d). Quantification of eRNA/lncRNA transcripts by smRNA-FISH (Fig. 4e) also revealed a similar trend to 3D contacts (Fig. 4c). Of note, recently annotated (https://www.ncbi.nlm.nih.gov/datasets/genome/GCF_000001635.27/) lncRNAs of unknown function overlap *Cd8* enhancers. We performed ChIP-seq for H3K27ac in CD69^-^DP progenitors^37^, CD4^+^8^lo^ differentiation intermediates, and CD4 and CD8 SP progeny and found that the developmental trajectory of H3K27ac levels broadly followed the same trend as 3D contacts (Extended Data Fig. 7d). Fig. 4f summarises the observed changes across modalities for CD4- and CD8-fated cells. These data reveal that the *Cd8* chromatin environment undergoes a complex series of developmental changes across multiple modalities, and chromatin contacts in particular show a striking pattern of transient attenuation.

### Transcriptional diversification occurs in an attenuated *Cd8* chromatin environment

Side-by-side comparison of chromatin contacts (Fig. 5a) and *Cd4/Cd8* transcriptional profiles (Fig. 5b) reveals that *Cd8* chromatin contact attenuation during CD4/CD8 lineage choice coincides with transcriptional diversification. It therefore appears that 3D contacts are selectively curtailed during a period of transcriptional exploration, creating a chromatin environment that remains transcriptionally permissive without enforcing high levels of *Cd8* expression. Interestingly, these findings are reminiscent of regulatory suboptimization, the concept that stronger is not always better in developmental gene regulation^42–46^, and that exploration of lineage options can benefit from attenuated interactions between transcription factors and developmental enhancers^42,43,45,46^ or preferential use of transcription factor paralogs of submaximal strength^44^. Hence, our data suggest that regulatory suboptimization may extend to 3D chromatin contacts.

**Figure 5.**
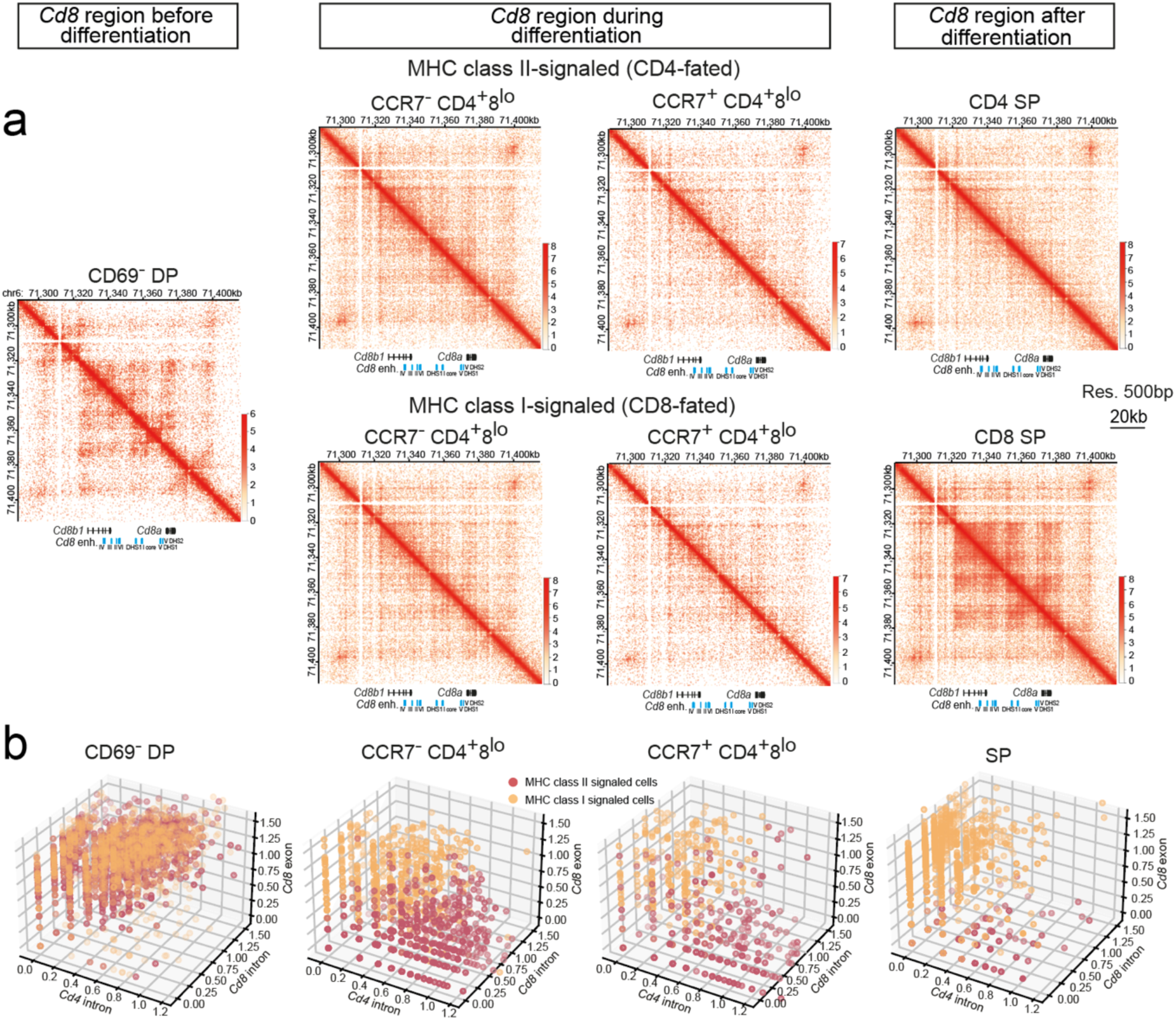
Transcriptional diversification occurs in an attenuated *Cd8* chromatin environment. Juxtaposition of chromatin contacts (a, from Fig. 4b) and *Cd4/Cd8* transcriptional profiles (b, from Fig. 2e). Note that the temporary attenuation of *Cd8* chromatin contacts during CD4/CD8 lineage choice coincides with the period of transcriptional diversification.

### Mesoscale contacts at the *Cd4* locus remain largely invariant during CD4/CD8 lineage choice and differentiation

We next focused on chromatin contacts at the *Cd4* locus (Fig. 6a). In contrast to the extensive, developmentally regulated 3D contact changes across the ∼60kb *Cd8* core region described above, inspection of RCPC data for *Cd4* revealed that mesoscale (10-100kb) contacts remained largely unchanged throughout the entire process of CD4/CD8 lineage choice and differentiation (Fig. 6b). This conclusion was supported by plotting observed over expected contacts (Extended Data Fig. 8) and quantification of chromatin loops (>5kb between loop anchors) called across the *Cd4* core region. Pair-wise comparison of the 70 loops identified in the *Cd4* core region between all 7 cell populations (n= 2940 comparisons) revealed only n=114 (3.88%) significant differences in loop strength (mean correlation R = 0.89). Strikingly, none of the 70 loops in the *Cd4* core region differed in strength between CD4 SP and CD8 SP progeny (Fig. 6c). The same comparison for the 82 loops identified in the *Cd8* core region (n= 3444 comparisons) showed 412 differential loops (11.96%, mean correlation R = 0.80, Fig. 6d). Hence, the *Cd4* core region undergoes limited mesoscale contact changes during CD4/CD8 lineage choice and differentiation compared to the *Cd8* core region.

**Figure 6.**
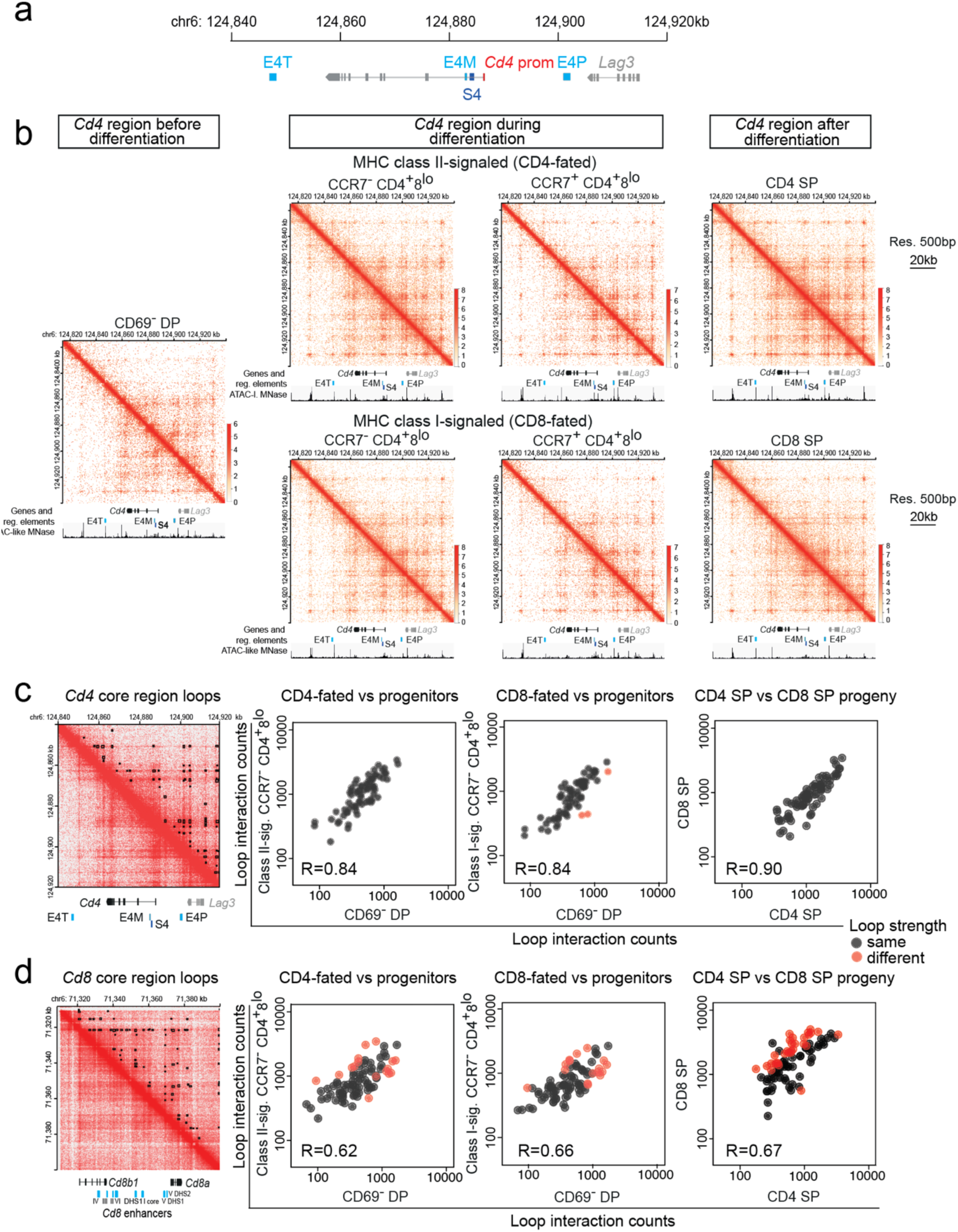
Mesoscale contacts at the *Cd4* locus remain largely invariant during CD4/CD8 lineage choice and differentiation. a, Schematic of the *Cd4* locus and its regulatory elements. *Cd4* prom: *Cd4* promoter, E4M: intronic enhancer, S4: intronic silencer, E4P: proximal enhancer, E4T: putative thymocyte enhancer^12^. b, RCPC maps (count matrix 500bp resolution, natural log scale), 2 merged biological replicates. ATAC-like tracks below the contact heatmaps show accessibility based on analysis of short-read MNase reads (Extended Data Fig. 7c). c, Left: the positions of chromatin loops with anchors at least 5kb apart and contained in the *Cd4* core region chr6:124840-124920kb are indicated in black in the top right half of the RCPC map (n=70). Log scale 0-5 (observed counts +1). Right: log/log loop interaction counts between the indicated thymocyte populations. Loops called as different in a particular comparison are marked in red. In total, 21 pair-wise comparisons of 70 loops between populations (n= 1470) by DeSeq2 showed n=57 (3.88%) significant differences and a mean correlation of R = 0.89. d, Left: the positions of chromatin loops with anchors at least 5kb apart and contained in the *Cd8* core region chr6:71310-71400kb are indicated in black in the top right half of the RCPC map (n=82). Log scale 0-5 (observed counts +1). Right: log/log loop interaction counts between the indicated thymocyte populations. Loops called as different in a particular comparison are marked in red. In total, 21 pair-wise comparisons of 82 loops between populations (n= 1722) by DeSeq2 showed n=206 (11.96%) significant differences and a mean correlation of R = 0.80.

### Master regulator transcription factors ThPOK and RUNX3 enforce CD4/CD8 lineage choice through a microscale *Cd4* enhancer/silencer switch

We used RCPC junction reads (Fig. 7a, see Methods) to visualise short-range chromatin contacts between the *Cd4* promoter and two closely spaced regulatory elements - the intronic *Cd4* enhancer and silencer (schematically depicted in Fig. 7b). The intronic *Cd4* silencer formed microscale contacts with the *Cd4* promoter in CCR7^-^ differentiation intermediates destined for both the CD4 and the CD8 lineages (Fig. 7c left, blue arrowhead). Contacts between the intronic *Cd4* enhancer and the *Cd4* promoter emerged selectively in CD4-fated CCR7^+^ differentiation intermediates (Fig. 7c center, red arrowhead). Of note, silencer-promoter contacts persisted in CD8-fated CCR7^+^ differentiation intermediates (Fig. 7c center, blue arrowhead). Enhancer-promoter contacts were further strengthened in mature CD4 SP cells (Fig. 7c right, red arrowhead). Fig. 7d shows the difference in reads between CD4 SP and CD8 SP. Red indicates contacts that are stronger in CD4 lineage cells, while blue indicates contacts that are stronger in CD8 lineage cells.

**Figure 7.**
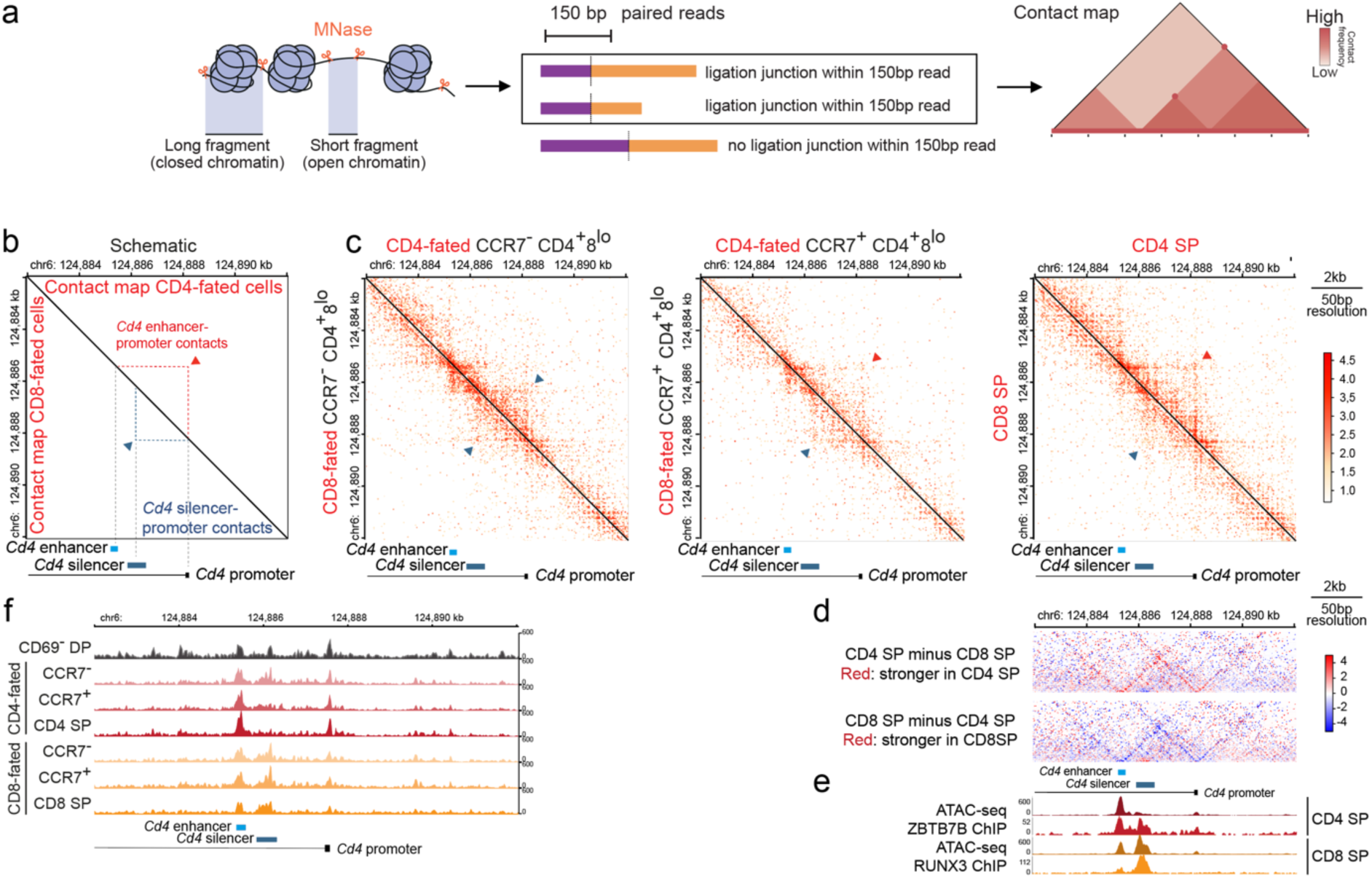
The T cell lineage master regulator transcription factors ThPOK and RUNX3 enforce lineage choice through a microscale enhancer/silencer switch at the *Cd4* locus. a, Use of MNase junction reads for contact map visualisation b, Schematic representation of predicted contacts between the Cd4 enhancer, silencer, and promoter c, RCPC junction reads at 50bp resolution, 2 merged biological replicates d, Subtraction plots of RCPC junction reads at 50bp resolution CD4 SP minus CD8 SP, 2 merged biological replicates. e, ATAC-seq and ZBTB7B ChIP-seq in CD4 SP (top) versus ATAC-seq accessibility and RUNX3 binding in CD8 SP (bottom). f, ATAC-like chromatin accessibility of the *Cd4* silencer is developmentally regulated. Note that the silencer is accessible in CD69^-^ progenitors and CCR7^-^ differentiation intermediates destined for both lineages and that accessibility selectively diminishes in CCR7^+^ differentiation intermediates destined for the CD4 lineage.

The CD4 lineage master transcription factor ThPOK/ZBTB7B is induced in CD4-fated cells at a late stage of CD4/CD8 T cell differentiation^16^ where it binds to *Cd4* enhancers^25^ (Fig. 7e, top) and antagonizes RUNX-mediated *Cd4* repression^27,28^ to enforce CD4 lineage fate^15^. Conversely, the CD8 lineage master transcription factor RUNX3 is induced in CD8-fated cells where it binds to and activates the *Cd4* silencer^26,29^ (Fig. 7e, bottom). Analysis of ATAC-like chromatin accessibility indicates the developmental regulation of *Cd4* silencer accessibility (Fig. 7f). The silencer is accessible in CD69^-^ progenitors and CCR7^-^ differentiation intermediates destined for both lineages, but accessibility begins to selectively diminish in CCR7^+^ differentiation intermediates destined for the CD4 lineage. Taken together, these data demonstrate how microscale contacts between the *Cd4* promoter and the *Cd4* enhancer or silencer are effectively ‘locked in’ following CD4/CD8 lineage choice.

### The role of regulatory suboptimization in CD4/CD8 lineage choice

To explore the importance of *Cd4* vs *Cd8* regulatory control, Singer and colleagues conceived transgenic mice in which cDNAs encoding *Cd8b1*-IRES-*Cd8a* and neomycin resistance were inserted into the 2nd exon of the *Cd4* gene^19^ (Extended Data Fig. 9a). The insertion is compact (∼2.5-3kb) and does not impinge on any of the known *Cd4* regulatory elements, with the intention of leaving the the remainder of the *Cd4* locus unperturbed (Extended Data Fig. 9a, b). We therefore used our RCPC data to explore the strength of contacts between regulatory elements driving transcription of *Cd8a/Cd8b1* from the *Cd4* promoter in this setting. We found that contacts between *Cd4* regulatory elements were less variable during CD4/CD8 lineage choice than contacts between *Cd8* regulatory elements at the native *Cd8* locus and did not undergo the transient attenuation described for the *Cd8* locus (compare contacts in Extended Data Fig. 9c to contacts in Fig. 4c). Interestingly, the consequence of placing *Cd8a/Cd8b1* into the *Cd4* promoter is a profound perturbation of CD4/CD8 lineage choice^19^. Perturbation of CD4/CD8 lineage choice was also described in transgenic mice expressing *Cd8a* under ectopic control^8,24^. Together, these data support the conclusion that transient reduction of *Cd8* expression is a critical element of CD4/CD8 lineage choice^3,4,17,19^ and our RCPC data provide new insights into the regulation of this process by 3D chromatin contacts.

## Discussion

Here we provide a single-molecule appraisal of CD4/CD8 lineage choice. We confirm that both MHC class I-signaled and MHC class II-signaled differentiation intermediates initially downregulate *Cd8* transcripts and provide direct evidence that MHC class I-signaled differentiation intermediates subsequently reverse this downregulation, consistent with coreceptor reversal^3,4,6,17^. While these data confirm key predictions of sequential (CD4-first) models of CD4/CD8 lineage choice^3,4,6^, single molecule analysis of lineage-defining transcripts pinpoints important quantitative differences. smRNA-FISH uncovers early bias in *Cd8a* transcript copy numbers imposed by TCR/co-receptor engagement via MHC class I versus MHC class II, which allows the identification of CD4- and CD8-fated differentiation intermediates by simple random forest classifiers. This observation is supported by qRT-PCR data for nascent and mature transcripts in sorted thymocyte populations performed in the current study and by published scRNA-seq data obtained by full-length Smart-seq2 cDNA sequencing^22^, which showed that *Cd8a* expression in MHC class I-signaled CD4^+^8^lo^ differentiation intermediates was several times higher than in MHC class I/II-signaled CD4^+^8^lo^ differentiation intermediates. Previous scRNA-seq studies reported variable results with respect to the emergence of transcriptional asymmetry. Chopp et al. reported moderately (∼50%) higher *Cd8a* transcript expression in *Ccr4^+^ Ccr7^-^* (’Sig1’) differentiation intermediates^21^, while Steier et al. found differences between MHC class I-versus MHC class II-signaled differentiation intermediates in the expression of signaling- and transcription factor-related transcripts, but not in *Cd8a* transcripts up to the point where late CD4^+^8^lo^ cells increase *Cd8a* expression indicative of coreceptor reversal^20^. We suggest that these observations are best explained by differences in the sensitivities of the assay systems employed^47^, which may underestimate transcriptional divergence.

The differences in *Cd8* downregulation between MHC class I- and II-signaled differentiation intermediates mean that *Cd4/Cd8* transcriptional divergence initiates earlier in CD4/CD8 lineage choice than anticipated by sequential models^3,4,6^. Transcriptional divergence also evolves more gradually than anticipated by one-step^7,9,10^ or sequential^3,4,6^ models, spanning both CCR7^-^ and CCR7^+^ CD4^+^8^lo^ stages of CD4/CD8 lineage choice. These findings raise interesting questions about the chromatin environment that underlies CD4/CD8 lineage choice.

To address these questions, we developed a low-input, ultra high-resolution region capture Pico-C approach, which we term RCPC. Our data reveal that the two lineage-defining loci, *Cd4* and *Cd8*, adopt fundamentally distinct strategies of 3D organisation to achieve developmental lineage- and stage-specific regulation. In bipotential progenitors prior to differentiation, the *Cd8* locus is organised by contacts that comprise the *Cd8a* and *Cd8b1* gene bodies and at least 6 developmentally regulated *Cd8* enhancers over the entire 60kb *Cd8* core locus. Differentiation intermediates of both lineages show transiently reduced 3D contacts across the entire *Cd8* locus. Transient attenuation of 3D contacts was accompanied by similar - albeit less strikingly coherent - changes in enhancer states and coding as well as non-coding transcription.

Transcriptional diversification occurs prior to the resolution of this temporary state of reduced mesoscale contacts at the *Cd8* locus, indicating that the transient attenuation of *Cd8* 3D contacts and other chromatin features provides a chromatin environment that is permissive for both ongoing transcription (in CD8-fated cells) and transcriptional silencing (in CD4-fated cells). Subsequently, mesoscale contacts at the *Cd8* locus are re-instated in CD8-fated cells in a reconfigured pattern consistent with the developmental state-specific activity of individual *Cd8* enhancer elements^14^, while residual contacts are lost in CD4-fated cells. By contrast, mesoscale contacts across the *Cd4* locus remain largely unchanged between progenitors, differentiation intermediates and differentiated progeny of both CD4 and CD8 lineages. At the microscale, however, contacts between the intronic *Cd4* enhancer and the *Cd4* promoter are lost in CD8-fated cells, and the intronic *Cd4* silencer, which is located between the *Cd4* enhancer and promoter is fully activated, as indicated by increased chromatin accessibility and prominent binding of the CD8 lineage-specific RUNX paralog RUNX3 (Ref.^29^).

The observation that transcriptional diversification is marked by transient weakening of *Cd8* locus-wide 3D contacts is reminiscent of regulatory suboptimization, the concept that stronger is not always better in developmental gene regulation, as previously described for developmental enhancers and transcription factor paralogs^42–46^. Regulatory suboptimization favours precision over brute force, facilitating appropriate transcriptional responses and ultimately lineage choice. To date, recognised manifestations of regulatory suboptimization act through transcription factor binding sites of submaximal strength in developmental enhancers^42,43,45,46^ and the preferential deployment of weaker transcription factor paralogs at critical developmental junctures^44^. Here we find that the *Cd8* locus undergoes transient attenuation of 3D contacts, which creates a permissive environment that is strong enough to support *Cd8* transcription, yet sufficiently malleable to allow for *Cd8* silencing. We propose that this form of ‘regulatory suboptimization in 3D’ facilitates the exploration of transcriptional states, thereby supporting appropriate T cell lineage choices. In summary, integration of quantitative transcript data and high-resolution analysis of 3D contacts reveals distinct regulatory strategies at lineage-defining loci.

## Acknowledgements

We thank Dr James Elliott and Bhavik Patel for cell sorting, Julia Merkenschlager for critical reading of the manuscript and Yuan Yin and Katie Croft (Baker lab) for discussion. Supported by the MRC (MM), the Wellcome Trust (099276/Z/12/Z to MM), the Simons Foundation (MP-SIP-00001828 to REB) and the Engineering and Physical Sciences Research Council (EP/Z534870/1 and EP/T517811/1 to REB).

## Methods

### Mice

Mouse work was done under a UK Home Office project licence and according to the Animals (Scientific Procedures) Act. C57BL/6 (C57BL/6OlaHsd, Envigo, UK) mice were used as wild-type controls. MHC I-deficient (JAX stock, 002070) and MHC II-deficient (JAX stock, 030434) mice were bred to generate MHC I, MHC II double-deficient mice. OT-I TCR transgenics (JAX stock, 003831) were bred with Rag1-deficient mice (MGI, 2448994). F5 TCR transgenics (MGI, 3706698) were bred with Rag1-deficient mice (JAX stock, 002216).

### Flow cytometry and Fluorescence-Activated Cell Sorting

Thymus and lymph nodes were dissected from 4-8 week old mice and single cell suspensions were stained with antibodies against CD4, CD8, CD69, TCR, CCR4, CCR7, CD71, NK1.1, B220 (BD Biosciences, eBioscience, Biolegend). Stained T cell populations were sorted using FACS Aria Fusion or FACS Aria III (BD Biosciences) or analysed using BD LSRFortessa or BD FACSymphony A3 (BD Biosciences). Flow cytometry FCS files were analysed with FlowJo v10 (TreeStar Inc).

### RNA extraction, reverse transcription and allele-specific qPCR

RNA was extracted from thymocytes using the RNeasy Mini Kit (Qiagen, 74106) according to manufacturer’s instructions, with DNase digestion for 30 min at RT. cDNA was synthesized using SuperScript III reverse transcriptase (ThermoFisher Scientific) following the manufacturer’s instructions. RT-qPCR was performed with QuantiTect Sybr Green PCR mix (Qiagen, 204145) and run on a CFX96 real-time PCR machine (Bio-Rad). Relative expression was calculated based on ΔCt values.

Primers

*Cd4* intron F (TCGCCTCTAGGCTTGTGTTT)

*Cd4* intron R (TAGGAGAGAGGCGATACGCA)

*Cd4* exon F (CCAGACAGTGTTCCTGGCTT)

*Cd4* exon R (TGCCTGGCGCTGTTGG)

*Cd8a* intron F (ACTGCTGATTTGGCTGCTCT)

*Cd8a* intron R (CTGCTCTTTCCACCCTCTGG)

*Cd8a* exon F (TACTACCAAGCCAGTGCTGC)

*Cd8a* exon R (ATCCGGTCCCCTTCACTGAG)

*UbcH7* F (TGGTTTGAATGGATACTCTGCTGGA)

*UbcH7* R (AGGAGGCTGATGAAGGAGCTTGA)

### smRNA-FISH

Sorted cells were seeded at a density of 3 – 10 x 10^5^ cells per well in an 18-well chamber slide (Ibidi, 81816) coated with poly-lysine. *Cd4*, *Cd8a*, *Actb* exon, intron and eRNA/lncRNA smRNA-FISH probes and View RNA ISH cell assay kit (QVC0001) were purchased from ThermoFisher and used according to the manufacturer instructions, with the following modifications: PBS was supplemented with 2nM Ribonucleoside-Vanadyl Complex and volumes were adjusted for 18-well chamber slide. Cells were imaged within a maximum of a week from staining using a Leica TCS SP5 inverted microscope with a 40x oil objective and a z-step size of 0.5 μm. For near-IR detection, Zeiss Axiovert 200M inverted fluorescence microscope was used with 60x oil objective.

### Image analysis and visualisation

Nuclei, cell outlines and dot identification were generated from maximum projections using CellProfiler v4.2.8. A size filter was applied to avoid scoring cell clusters as individual cells. Outlines have been checked manually for accuracy, and cells visibly damaged or dividing were eliminated. Data were plotted using matplotlib v3.4.3. Co-localisation analysis was done in ImageJ. For each image, an area was created from the dots selected, and the overlap was calculated in ROI Manager using AND command. The percentage of overlap was calculated by dividing the AND area by the intronic area.

The distribution of cells in gene-expression space based on *Cd4* intron, *Cd8* intron/exon was analysed using kernel density estimators. For both class I- and II-signaled cells, a density field was computed using a shared grid, and an isosurface was extracted at the 90 percent threshold. The volume of the isosurface was used as a quantitative measure of the spread of cells in gene-expression space. Pairwise comparisons between MHC class I- and MHC class II-signaled populations were then performed by calculating the dice overlap score (using D = 2 x volume(A intersect B)/(volume(A) + volume(B)), where A and B are the regions enclosed by the two isosurfaces) between the MHC class I- and MHC class II-signaled populations. To compare subsequent stages during development, the KDE volumes were normalised to the CD69-DP population as a common reference point.

### Transcriptional burst identification and burst fraction quantification

Intron signals were identified by automatic dot detection using CellProfiler and quantified as a sum of the dot mean radius and intensity of the brightest pixel (Max Intensity). The threshold for defining a burst was set at the base of the upper size distribution, corresponding to the point at which larger objects become distinct from the main population. The size threshold was established on a per-experiment basis due to experimental variability (microscope acquisition settings, fluorophore characteristics, and hybridisation efficiency). Threshold values generally ranged between 2.0 and 2.5, and where necessary, the selected threshold was validated using positive and negative control samples.

### Random Forest Classifier

We trained a random forest classifier developed using scikit-learn in Python 3 on an 80/20 train/test split of single-cell data (n = 7,739) comprising seven fine-grained population labels (CD69⁻ DP, MHC I-or II-signaled CCR4⁺7⁻, MHC I-or II-signaled CCR7⁺, CD4 SP, or CD8 SP) and five smRNA-FISH measurements (*Cd8a* exon count, *Cd8a* intron count, *Cd8a* transcriptional burst count, *Cd4* intron count, and *Cd4* transcriptional burst count, cells missing any of these were removed). Stratified sampling was used to preserve the proportions of each population label in the train and test sets. Hyperparameters were selected using a randomised search (n=40 iterations), scored by weighted F1 using five-fold stratified cross-validation with three repeats. In testing, fine-grained populations were aggregated into i) progenitors, ii) CD4 or iii) CD8 and the classifier was used to predict which fine-grained population and aggregated population each cell in the test data belonged to. Feature importance was impurity-based. For classification of six sorted F5 (n = 735) and OT-I (n = 12,489) subsets, all data in the original 80/20 train/test split were used to train the classifier.

### RCPC

Thymocytes were sorted as follows: CD69-DP (1 thymus/ replicate, MHC class I/II double KO CD69^-^CD71^-^CD4^+^CD8^+^), MHC II-signaled CCR7^-^ (2 thymi/replicate, MHC class I KO CD69^+^CCR7^-^CD4^+^CD8^lo^), MHC II-signaled CCR7^+^ (2 thymi/replicate, MHC class I KO CD69^+^CCR7^+^CD4^+^CD8^lo^), CD4 SP (2 thymi/replicate, MHC class I KO CD69^-^CD4^+^CD8^-^ TCR^+^), MHC I-signaled CCR7^-^ (4 thymi/replicate, MHC class II KO CD69^+^CCR7^-^ CD4^+^CD8^lo^NK1.1^-^), MHC I-signaled CCR7^+^ (4 thymi/replicate, MHC class II KO CD69^+^CCR7^+^CD4^+^CD8^lo^NK1.1^-^), CD8 SP (4 thymi/replicate, MHC class II KO CD69^-^CD4^-^ CD8^+^TCR^+^). Thymocytes were simultaneously crosslinked with 3 mM EGS (ethylene glycol bis(succinimidyl succinate)) for a total of 45 min at RT and 1% formaldehyde added for the last 15 minutes. The reaction was quenched using 1.2 M Tris pH 7.5 for 5 min. Cells were centrifuged, and the cell pellet was snap-frozen in liquid nitrogen and stored at -80°C. Pico-C libraries were generated following the published protocol^34^, available at protocols.io (https://doi.org/10.17504/protocols.io.kqdg31nm1l25/v1). Chromatin was digested with 7.5 U MNase (micrococcal nuclease) for 10 min at 37°C with 950 rpm shaking. To assess the number of cycles needed for the library preparation, 20% of the sample was used for pilot PCR, the samples were run for 11-13 cycles depending on the initial DNA content.

A Custom Twist panel was designed to capture interactions with loci of interest *Cd4* chr6: 124,702,023 – 125,100,595 and *Cd8* chr6: 71,033,496 – 71,762,124 (mm10). Up to 8 indexed libraries were pooled and the DNA was dried using a vacuum concentrator at 35°C for 2 hours. The capture was performed according to the Twist Target Enrichment Standard Hybridisation v1 Protocol and as described^35^ with the following modifications: Mouse COT-1 DNA blocker solution was used instead of Twist’s Blocker Solution, and the libraries were divided in two and run for 11 PCR cycles. DNA was quantified using Qubit (Invitrogen) and sent for sequencing to Novogene (sequencing platform NovaSeq X Plus Series PE150).

### RCPC data analysis

FASTQ files of paired end reads for each sample/lane were downloaded from NovoGene. FASTQ file quality was confirmed using FastQC v0.12.1, and reads were aligned to the GRCm38 (mm10) mouse genome using BWA-MEM v0.7.19 with - T0 -5SP. Aligned reads were processed to contact pairs using pairtools v1.1.3 (https://doi.org/10.1371/journal.pcbi.1012164) with parse2 --min-mapq 3 --report-position outer --max-inter-align-gap 30 --max-insert-size 250 --add-pair-index --add-columns pos5,pos3, and deduplicated with pairtools dedup --max-mismatch 1. Replicates for each stage were pooled with pairtools merge and then filtered and separated into separate files that included either the *Cd4* or *Cd8* capture loci with pairtools select. Pairs were converted to contact matrices in cooler (Ref.^48^) and Juicer (Ref.^49^) formats with cooler cload pairs v0.10.2 and juicer tools pre v2.20 for resolutions between 50bp-10kb. Contact maps were balanced by ICE normalisation using cooler balance. Loops were called using chromosight detect (Ref. ^50^) with flags --min-dist 5000 -i 3 -P loops -z 80 -n raw --min-separation 250 at resolutions 250bp and 500bp for CD4 SP and CD8 SP maps. Loop calls were clustered and aggregated using the clusterPairs function of the bioconductor package diffHic (Ref.^51^) with tol = 200, upper = 2000. The central pixels from each aggregated loop were extracted for each replicate and interaction counts compared across stages using DeSeq2 to determine significantly changing interactions for each pairwise comparison.

### Junction read analysis

Analysis of ligation junction positioning was processed similarly to outer positions but with the pairtools parse2 flag --report-position junction. Pairs with a detectable ligation junction position were extracted based on the index tag reported by pairtools, with exact positions recoverable for ∼40% of reads on average. Exact lengths of each fragment end of this subset were determined by comparing read outer vs junction positions. Where the ligation site was sequenced on only read 1 or read 2, the longer sub-fragment was assumed to be contiguous if the genomic distance between the 5’ and 3’ end was <31bp.

For short-fragment chromatin accessibility analysis, sub-70bp fragment ends were converted to bampe format using pysam. Peaks were called with MACS3 in bampe mode with -q 0.01. Reproducible peaks between biological replicates were pooled across stages, and deseq2 was used to compute scale factors on the basis that peak coverage should be consistent between stages on the scale of the entire ∼1.1Mb of both capture regions.

### Contact Strength comparisons

Regions around Cd8 enhancers and promoters were defined manually based on inspection of RCPC contact maps. Balanced Interactions were extracted at 250bp resolution using the Bioconductor package HiCExperiment (Ref.^52^) and compared using ggplot2. For comparisons in Extended Data 9, contacts were normalised against the expected value at each distance, due to the close proximity between features. Expected interactions were calculated using CoolTools (Ref.^53^).

### MNase-based chromatin accessibility

The RCPC design uses 150bp paired-end sequencing and size selection for fragments of approximately dinucleosomal length and results in ∼40% of reads that contain a sequenced ligation junction. As noted previously^41^ this provides the precise length and position of each interacting fragment end comprising the contact pairs. We noted that ends of subnucleosomal length were enriched at open chromatin and used junction read fragments <70bp as a proxy for MNase-based chromatin accessibility.

### RCPC junction reads for short-range contacts

At very short ranges, non-junction reads contain less informative on-diagonal interactions (likely proximal nucleosome-nucleosome interactions) while junction reads are enriched for contacts between open chromatin.

### ChIP-seq, ATAC-seq and other genomic features

ChIP-seq data was generated as previously described^37^ by sequencing 50 bp single-ended reads on Illumina HiSeq 2500. CD69^−^ DP and CD4 SP CTCF and RAD21 ChIP-seq data are available in GSE199059. Read quality and preprocessing were checked with Fastp v0.23.4 (Ref.^54^). Reads were mapped to the mm10 genome with Bowtie 2 v2.5.3 (Ref.^55^) and deduplicated with samtools. Coverage tracks were produced using BedTools v2.31.1 (Ref.^56^) at a binsize of 50 bp.

H3K27ac ChIP-seq in CD69-DP, CD4^+^8^lo^ differentiation intermediates, and CD4 and CD8 SP was similarly sequenced and processed. Values in heatmaps quantifying H3K27ac signal at Cd8 enhancers were calculated with deseq2 vst, merging replicates and scaling relative to the max at each enhancer.

Contact domains, compartments and subcompartments were defined as described in Guo et al. 2022 on CD69-DP Hi-C and converted from mm9 to mm10 coordinates with liftOver (doi.org/10.1101/gr.229102). Processed Ring1B ChIP-on-chip was obtained from GSM1298086 and converted from mm8 to mm10 using liftOver; tracks display logFold2 signal over control.

For analysis of CD8 SP Runx3 ChIP data^29^ (GSE280658) and CD4 SP ZBTB7B ChIP data^21^ (GSE148976), reads were trimmed with trim_galore (doi:10.5281/zenodo.5127899), aligned with bowtie2, and deduplicated with samtools. RPGC-normalised coverage tracks were produced with deepTools bamCoverage (doi:10.1093/nar/gkw257). ATAC-seq data for CD69^-^ DP, CD4 SP and CD8 SP tracks were downloaded from GSE107076. scRNA-seq data^22^ were downloaded from GSE149207.

### Data Visualisation

Genome browser snapshots for visualising contact maps, RNA-seq, scRNA-seq, ChIP-seq and ATAC-seq were produced using CoolBox (v0.3.9) (Ref.^57^), JuiceBox version 2.17.00 (Ref.^49^) or the IGV Integrative Genomics Viewer (v2.12.0) (Ref.^58^). Contact maps were plotted with raw rather than balanced counts, as manual inspection determined that balancing contact matrices resulted in a loss of clarity of loop and stripe structure at higher resolutions.

## Data availability

High throughput sequencing data generated for this study will be deposited at GEO. H3K27ac, RAD21, CTCF and H3K27me3 ChIP-seq data^37^ and Hi-C data^37^ used to call compartments and subcompartments are available in the GEO database under accession code GSE199059. RING1B ChIP-on-chip data^29^ are available in the GEO database under accession code GSM1298086. RUNX3 ChIP data^29^ are available in the GEO database under accession code GSE280658. ZBTB7B ChIP data^21^ are available in the GEO database under accession code GSE148976. ATAC-seq data^38^ are available in the GEO database under accession code GSE107076. Full-length scRNA-seq data^22^ are available in the GEO database under accession code GSE149207. Source data are provided with this paper.

## Code availability

Code generated for this study will be available on Github.

---

**Supplementary Figure 1.**
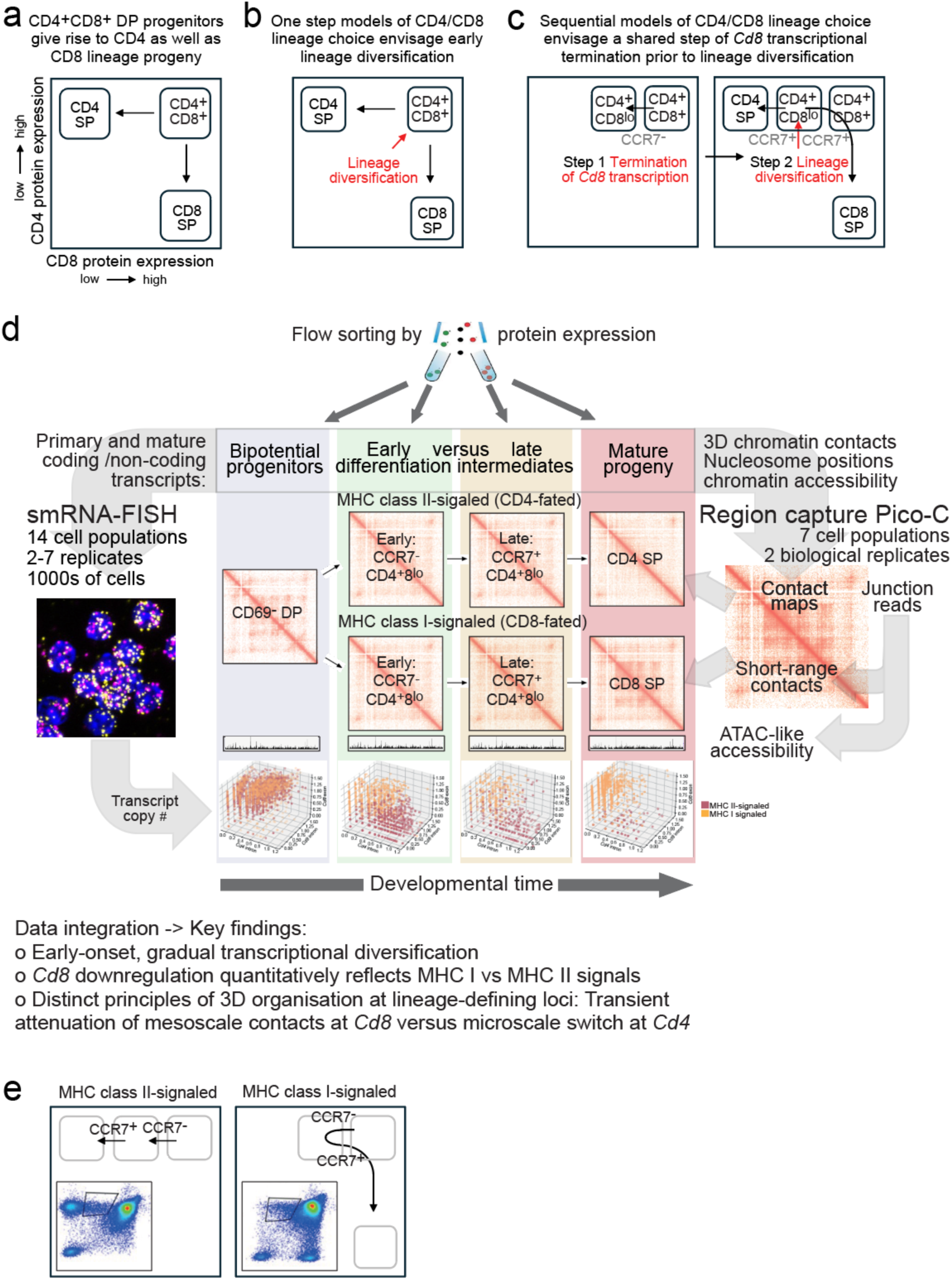
Models of CD4/CD8 lineage choice and study overview. a, CD4 helper and CD8 cytotoxic T cells arise from a shared bipotential progenitor b, One step models of CD4/CD8 lineage choice c, Sequential (CD4 first) models of CD4/CD8 lineage choice d, Study overview e, Schematic of thymocyte differentiation trajectories in mice that selectively express either MHC class I or MHC class II. Insets show illustrative flow cytometry plots of CD4 and CD8 expression.

**Extended Data Figure 1.**
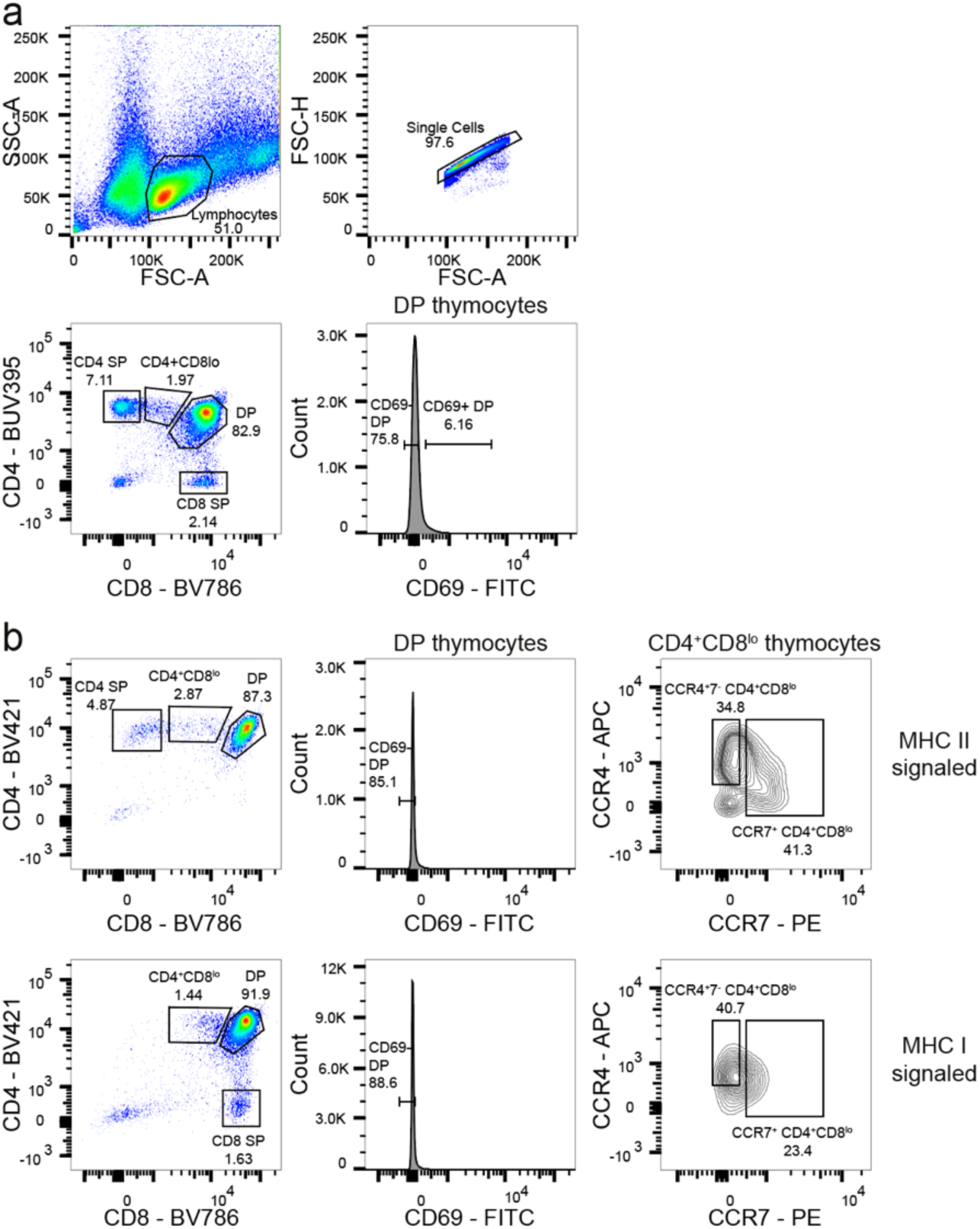
Gates for flow cytometry. a, Sort gates for Fig. 1 b, Sort gates for Fig. 2 and subsequent figures, except Extended Data Fig. 4, which shows its own gating strategy

**Extended Data Figure 2.**
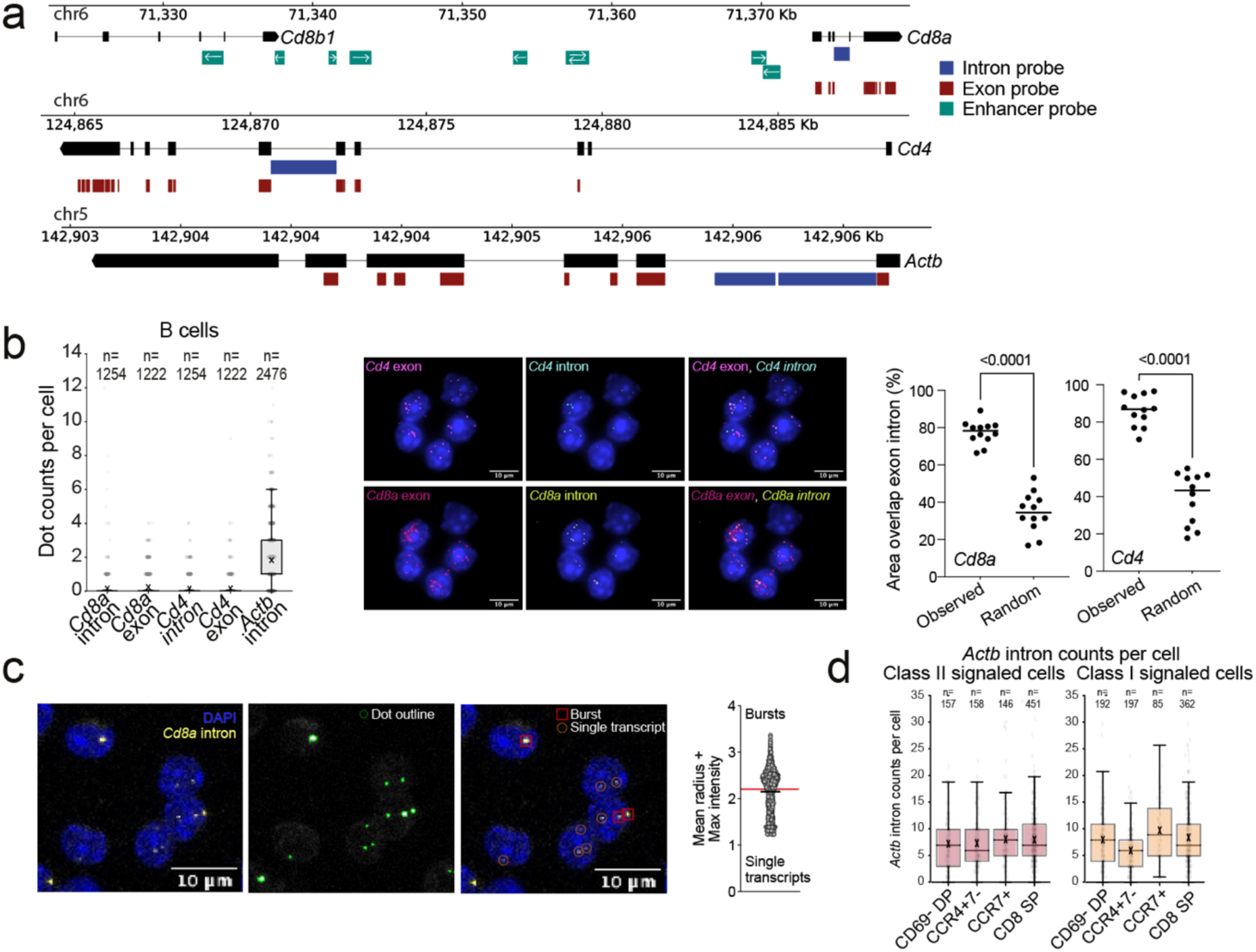
smRNA-FISH probes. a, Genomic positions of exon, intron and lnc/eRNA-FISH probes. Exon probes were pre-designed by the manufacturer. Intron and lnc/enh probes were designed to hybridise with sequences of interest. Horizontal bars indicate probe position and arrows indicate the directionality of lncRNA/enhancer RNA probes. b, Probe specificity was assessed by testing *Cd4* and *Cd8a* probes in B cells (left, 2 biological replicates) and by assessing the specific overlap between *Cd4* intron/exon probes and *Cd8a* intron/exon probes in thymocytes (right, 3 biological replicates). Random: overlap between *Cd4* intron/ *Cd8a* exon probes and *Cd8a* intron/*Cd4* exon probes. Stats: unpaired t-test. c, Definition of transcriptional bursts in smRNA-FISH images (left image). Intron signals were identified by automatic dot detection using CellProfiler (center image) and quantified as a sum of the dot mean radius and intensity of the brightest pixel (Max Intensity) as illustrated in the graph. Example bursts and single transcripts are shown (right image). See methods for details. d, Overall transcription activity was assessed using intron probes for the housekeeping gene *Actb.* n = the number of cells scored for each probe and cell population, 1 biological replicate.

**Extended Data Figure 3.**
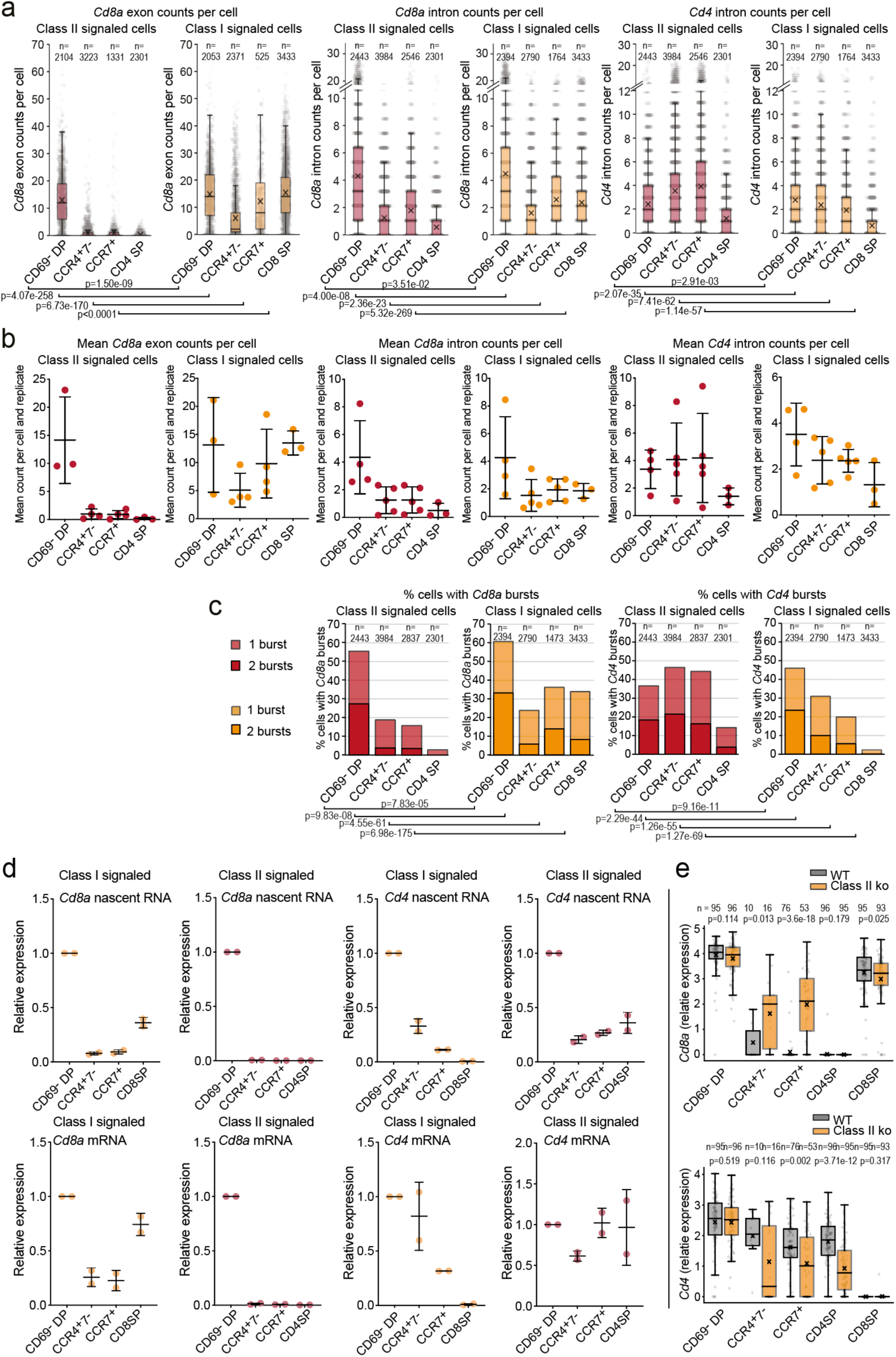
smRNA-FISH counts, qRT-PCR and Smart-seq2 data. a, *Cd8a* exon counts (left, from Fig. 2c), *Cd8a* intron counts (middle), and *Cd4* intron counts (right) determined by smRNA-FISH for the indicated cell populations. n = number of cells scored for each probe and cell population. Stats: Chi square test. b, Mean counts per replicate for the data shown in b. c, Percentage of cells within the indicated subsets that show transcriptional bursts for *Cd8a* (left) and *Cd4* (right). From Fig. 2b. Stats: Chi square test. d, qRT-PCR of the indicated transcripts. 2 biological replicates. Expression is normalised to *UbcH7* and shown relative to CD69-DP e, scRNA-seq data showing *Cd4* and *Cd8a* expression in the indicated subsets of MHC class I-signaled versus MHC class I/II-signaled CD4^+^8^lo^ thymocytes as determined by full-length cDNA sequencing^22^. Stats: Mann-Whitney U test.

**Extended Data Figure 4.**
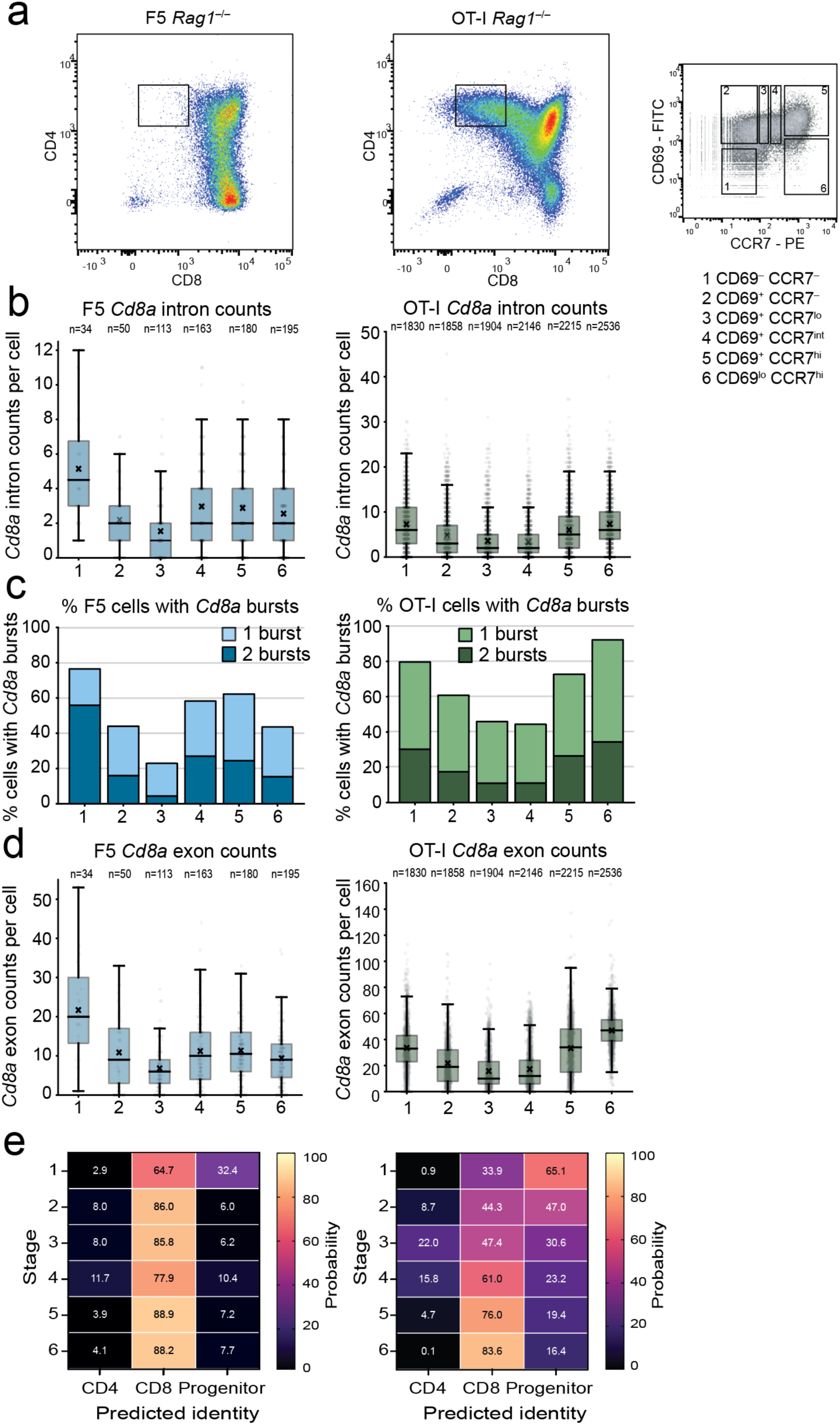
Transcriptional downregulation of *Cd8a* in MHC class I-signaled TCR transgenic differentiation intermediates. a, Isolation of differentiation intermediates as well as pre- and post-differentiation populations from F5 (left) and OT-I (right) TCR transgenic thymus according to the Kimura et al., 2016 scheme^18^ (far right). Live and single cell gates as shown in Extended Data Fig. 1a. Stages 1 and 6 were further enriched by gating on CD4 and CD8 (see Extended Data Fig. 1a, b). b, *Cd8a* intron counts determined by smRNA-FISH. 1 biological replicate for F5 and 2 biological replicates for OT-I. c, *Cd8a* transcriptional bursts determined by smRNA-FISH d, *Cd8a* exon counts determined by smRNA-FISH. Note that F5 TCR transgenic differentiation intermediates, which do not form a prominent CD4^+^8^lo^ population of differentiation intermediates, nevertheless transiently downregulate *Cd8a* transcripts. e, Random forest classification of sequential developmental stages in F5 and OT-I TCR transgenic thymocytes.

**Extended Data Figure 5.**
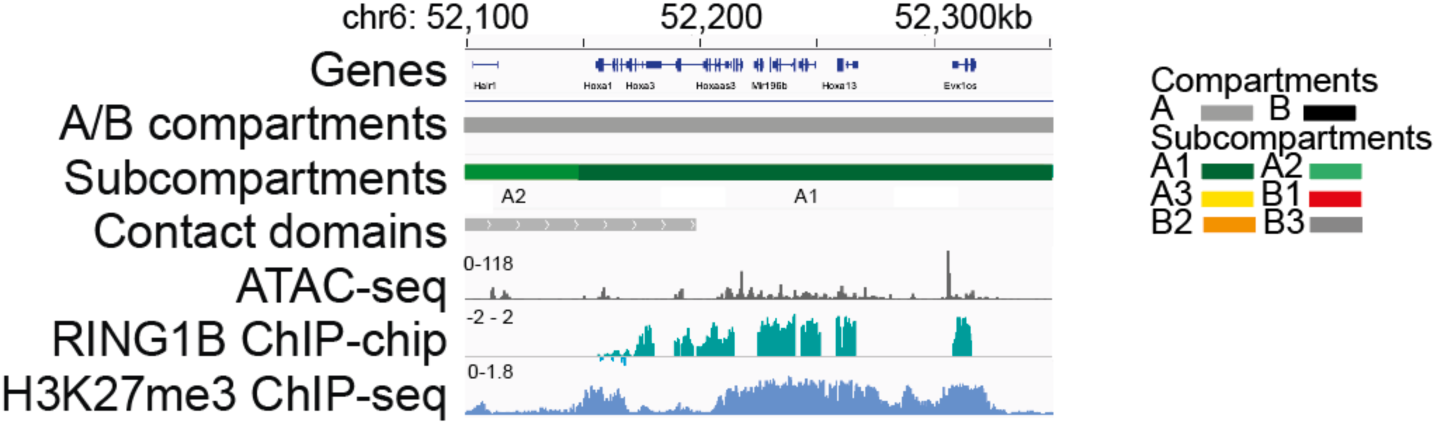
Genome browser view of the HoxA cluster on mouse chromosome 6. This region serves as a positive control for Fig. 3d, demonstrating the presence of the PRC1 component RING1B and the PRC2-catalysed repressive histone mark H3K27me3 in developing T cells.

**Extended Data Figure 6.**
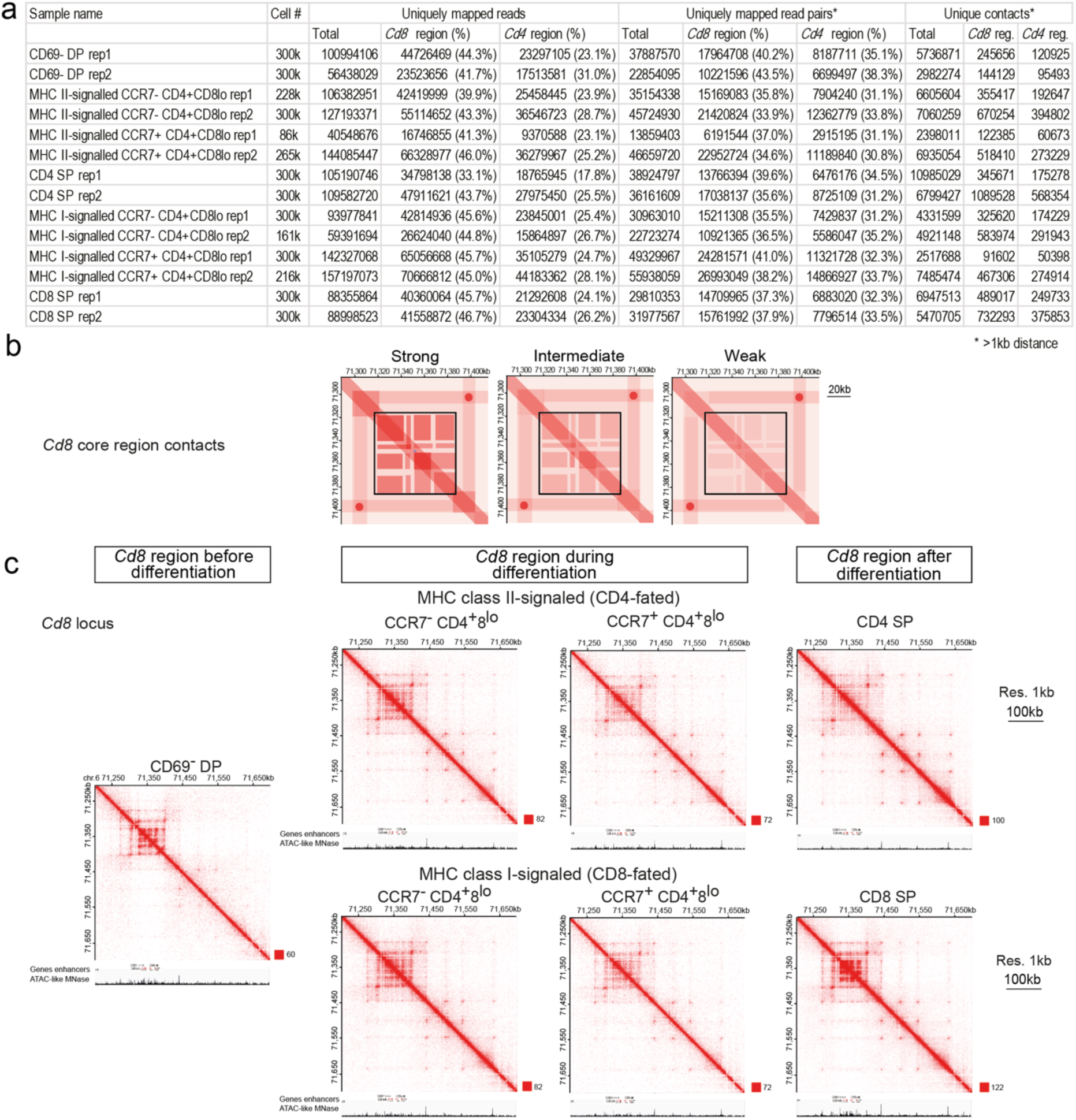
Attenuation of 3D contacts is selective for the *Cd8* core region. a, Summary of RCPC experiments. Numbers and percentages of uniquely mapped reads, read pairs, and contacts. See methods for sort strategies. Sig.: signalled, R: replicate. b, Schematic depiction of key changes in chromatin conformation data shown in Fig. 4b. A black outline marks the *Cd8* core region c, A broader view of the *Cd8* capture region shows persistence of microcompartments and other fine-scale features across the *Cd8* capture region during CD4/CD8 lineage choice and differentiation, which indicates that the temporary attenuation of 3D contacts described in Fig. 4b is selective for gene-enhancer contacts in the *Cd8* core region.

**Extended Data Figure 7.**
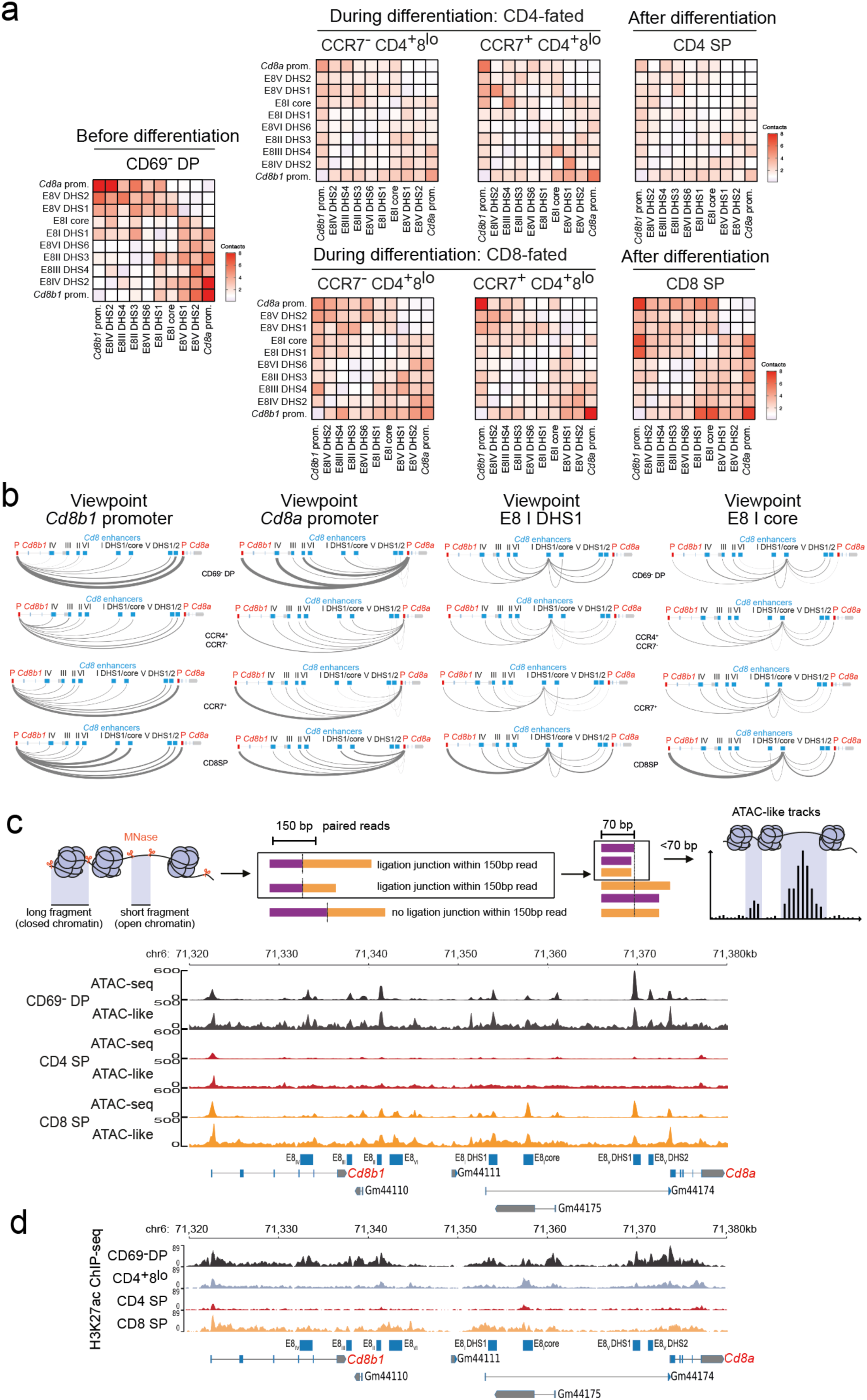
Developmental dynamics of chromatin contacts, accessibility and H3K27ac at *Cd8* regulatory elements. a, Developmental regulation of observed/expected contacts made by each *Cd8* regulatory element with each other *Cd8* regulatory element as determined by RCPC. b, Developmental reconfiguration of observed over expected RCPC contacts shown from the perspective of the *Cd8b1* promoter, the *Cd8a* promoter, *Cd8* enhancer I DHS1 (E8 I DHS1) and *Cd8* enhancer I core (E8 I core) in CD8-fated cells. c, As detailed in Methods, ∼40% of paired end 150bp RCPC reads contain a ligation junction. The shorter fragments of these junction reads are enriched at open chromatin, similar to previous observations of short MNase fragment distributions^40,41^. Note overlap of peak positions between ATAC-seq and MNase accessibility tracks. d, Developmental regulation of H3K27ac levels at *Cd8* regulatory elements. H3K27ac ChIP-seq was performed for the indicated cell populations. One of 2 biological replicates.

**Extended Data Figure 8.**
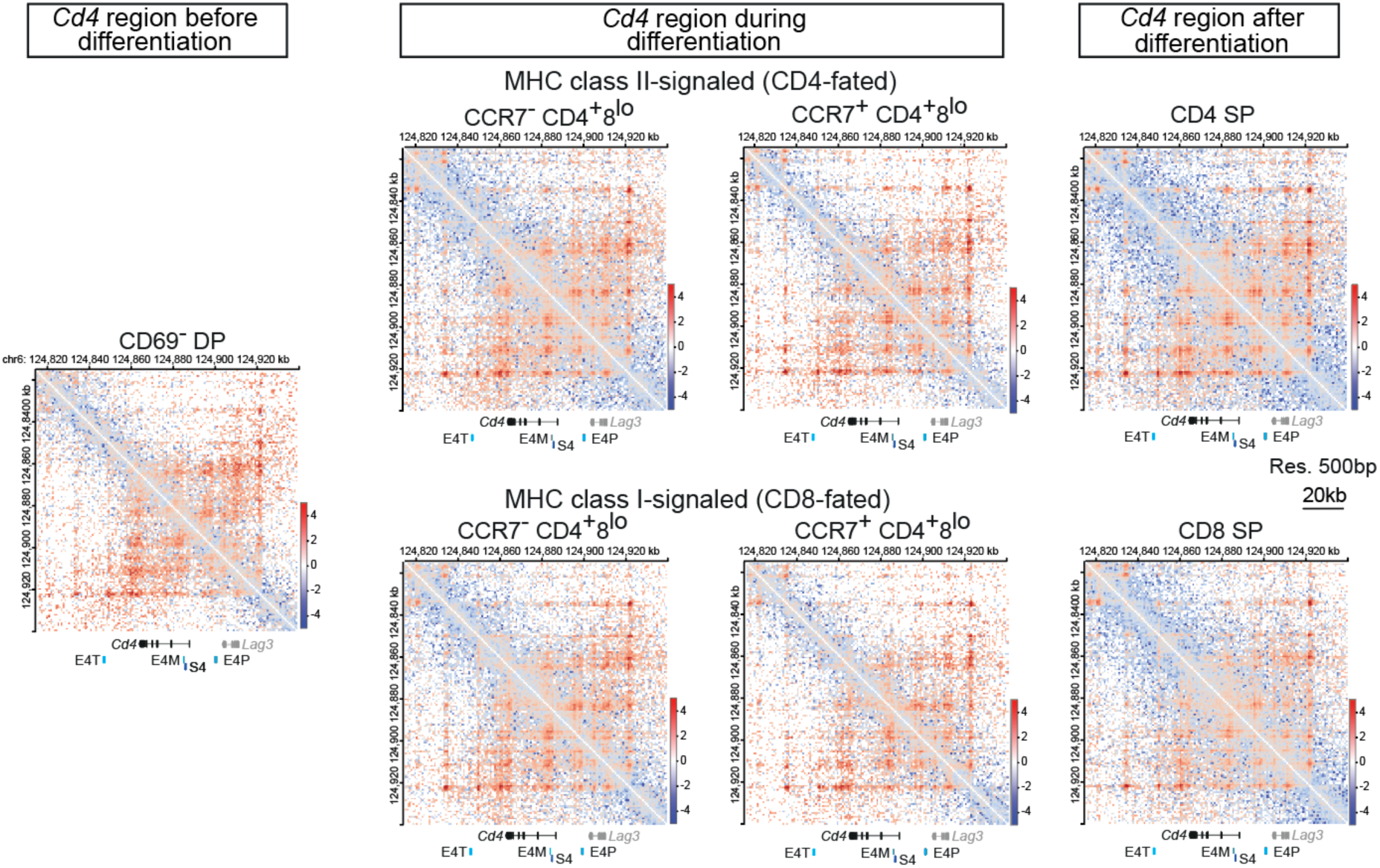
Observed over expected contacts and loop quantification at the *Cd4* locus. Observed over expected contacts at the *Cd4* locus before, during and after CD4/CD8 lineage choice and differentiation at 500bp resolution.

**Extended Data Figure 9.**
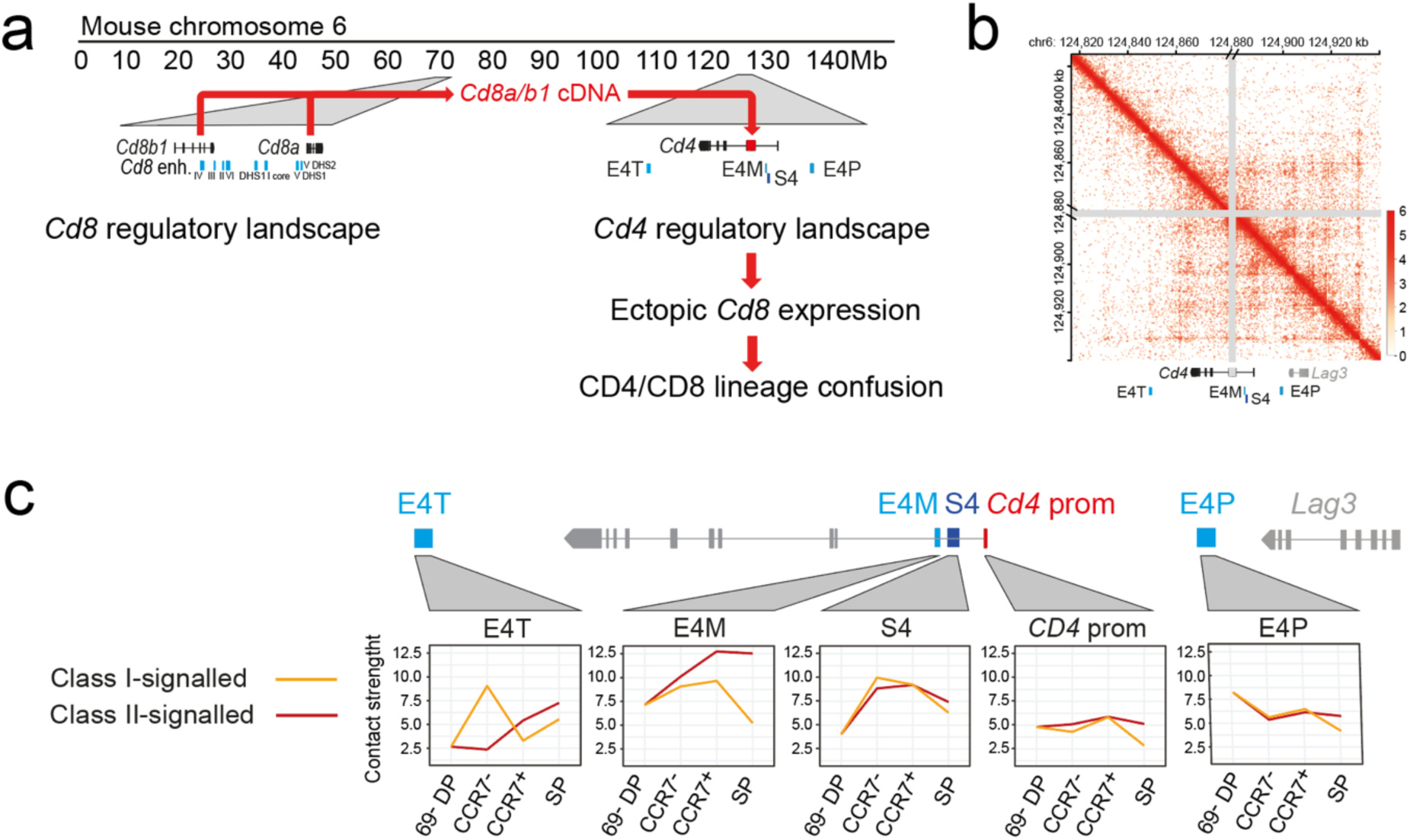
*Cd8* transgene expression from the *Cd4* locus perturbs CD4/CD8 lineage choice - predicted impact on developmentally regulated 3D contacts. a, Experimental outline b, RCPC map of the *Cd4* locus with insertion of a *Cd8a/b1/NeoR* cDNA construct into the 2nd exon of *Cd4* (Ref.^19^) indicated in grey. Count matrix 500bp resolution, natural log scale. c, Observed over expected contacts between the *Cd4* promoter, the intronic enhancer E4M and silencer S4, the proximal enhancer E4P, and the putative thymocyte enhancer E4T (Ref.^12^). Note that the *Cd8a/b* transgene insertion leaves the spatial arrangement between E4P, E4M, S4, and E4P unchanged, and only slightly increases the distance between E4P and E4T.

## References

1. von Boehmer, H. The developmental biology of T lymphocytes. Annu. Rev. Immunol. 6, 309–326 (1988).

2. Germain, R. N. T-cell development and the CD4-CD8 lineage decision. Nat. Rev. Immunol. 2, 309–322 (2002).

3. Singer, A. New perspectives on a developmental dilemma: the kinetic signaling model and the importance of signal duration for the CD4/CD8 lineage decision. Curr. Opin. Immunol. 14, 207–215 (2002).

4. Singer, A., Adoro, S. & Park, J. H. Lineage fate and intense debate: myths, models and mechanisms of CD4-versus CD8-lineage choice. Nat. Rev. Immunol. 8, 788–801 (2008).

5. Taniuchi, I. CD4 helper and CD8 cytotoxic T cell differentiation. Annu. Rev. Immunol. 36, 579–601 (2018).

6. Steier, Z., Kim, E. J. Y., Aylard, D. & Robey, E. A. The CD4 Versus CD8 T Cell Fate Decision: A Multiomics-Informed Perspective. Annu. Rev. Immunol. 42, 235–258 (2024).

7. Robey, E. A. et al. Thymic selection in CD8 transgenic mice supports an instructive model for commitment to a CD4 or CD8 lineage. Cell 64, 99–107 (1991).

8. Robey, E., Itano, A., Fanslow, W. C. & Fowlkes, B. J. Constitutive CD8 expression allows inefficient maturation of CD4+ helper T cells in class II major histocompatibility complex mutant mice. J. Exp. Med. 179, 1997–2004 (1994).

9. Chan, S. H. et al. Another view of the selective model of thymocyte selection. Cell 73, 225–236 (1993).

10. Davis, C. B. et al. Evidence for a stochastic mechanism in the differentiation of mature subsets of T lymphocytes. Cell 73, 237–247 (1993).

11. Sawada, S., Scarborough, J. D., Killeen, N. & Littman, D. R. A lineage-specific transcriptional silencer regulates CD4 gene expression during T lymphocyte development. Cell 77, 917–929 (1994).

12. Adlam, M. & Siu, G. Hierarchical interactions control CD4 gene expression during thymocyte development. Immunity 18, 173–184 (2003).

13. Siu, G. et al. A transcriptional silencer controls the developmental expression of the CD4 gene. EMBO J. 13, 3570–3579 (1994).

14. Ellmeier, W., Sawada, S. & Littman, D. R. The regulation of CD4 and CD8 coreceptor gene expression during T cell development. Annu. Rev. Immunol. 17, 523–554 (1999).

15. Kappes, D. J., He, X. & He, X. Role of the transcription factor Th-POK in CD4:CD8 lineage commitment. Immunol. Rev. 209, 237–252 (2006).

16. Egawa, T. & Littman, D. R. ThPOK acts late in specification of the helper T cell lineage and suppresses Runx-mediated commitment to the cytotoxic T cell lineage. Nat. Immunol. 9, 1131–1139 (2008).

17. Brugnera, E. et al. Coreceptor reversal in the thymus: signaled CD4+8+ thymocytes initially terminate CD8 transcription even when differentiating into CD8+ T cells. Immunity 13, 59–71 (2000).

18. Kimura, M. Y. et al. Timing and duration of MHC I positive selection signals are adjusted in the thymus to prevent lineage errors. Nat. Immunol. 17, 1415–1423 (2016).

19. Shinzawa, M. et al. Reversal of the T cell immune system reveals the molecular basis for T cell lineage determination in the thymus. Nat. Immunol. 23, 731–742 (2022).

20. Steier, Z. et al. Single-cell multiomic analysis of thymocyte development reveals drivers of CD4+ T cell and CD8+ T cell lineage commitment. Nat. Immunol. 24, 1579–1590 (2023).

21. Chopp, L. B. et al. An integrated epigenomic and transcriptomic map of mouse and human αβ T cell development. Immunity 53, 1182–1201.e8 (2020).

22. Karimi, M. M. et al. The order and logic of CD4 versus CD8 lineage choice and differentiation in mouse thymus. Nat. Commun. 12, 99 (2021).

23. Yayon, N. et al. A spatial human thymus cell atlas mapped to a continuous tissue axis. Nature 635, 708–718 (2024).

24. Itano, A. et al. The cytoplasmic domain of CD4 promotes the development of CD4 lineage T cells. J. Exp. Med. 183, 731–741 (1996).

25. Kojo, S., Yasmin, N., Muroi, S., Tenno, M. & Taniuchi, I. Runx-dependent and silencer-independent repression of a maturation enhancer in the Cd4 gene. Nat. Commun. 9, 3593 (2018).

26. Taniuchi, I. et al. Differential requirements for Runx proteins in CD4 repression and epigenetic silencing during T lymphocyte development. Cell 111, 621–633 (2002).

27. Wildt, K. F. et al. The transcription factor Zbtb7b promotes CD4 expression by antagonizing Runx-mediated activation of the CD4 silencer. J. Immunol. 179, 4405–4414 (2007).

28. Muroi, S. et al. Cascading suppression of transcriptional silencers by ThPOK seals helper T cell fate. Nat. Immunol. 9, 1113–1121 (2008).

29. Ogawa, C. et al. Phosphorylation of Runx protein controls helper CD4+ T cell versus cytotoxic CD8+ T cell lineage choice. Nat. Immunol. 27, 799–811 (2026).

30. Teschendorff, A. E. & Enver, T. Single-cell entropy for accurate estimation of differentiation potency from a cell’s transcriptome. Nat. Commun. 8, 15599 (2017).

31. Dussiau, C. et al. Hematopoietic differentiation is characterized by a transient peak of entropy at a single-cell level. BMC Biol. 20, 60 (2022).

32. Ehrlich, L. I. R., Oh, D. Y., Weissman, I. L. & Lewis, R. S. Differential contribution of chemotaxis and substrate restriction to segregation of immature and mature thymocytes. Immunity 31, 986–998 (2009).

33. Li, Y. et al. CCR4 and CCR7 differentially regulate thymocyte localization with distinct outcomes for central tolerance. eLife 12, e80443 (2023).

34. Maziak, N. et al. Three-dimensional genome reorganization foreshadows zygotic genome activation in Drosophila. Nat. Genet. 58, 607–617 (2026).

35. Goel, V. Y., Huseyin, M. K. & Hansen, A. S. Region Capture Micro-C reveals coalescence of enhancers and promoters into nested microcompartments. Nat. Genet. 55, 1048–1056 (2023).

36. Hsieh, T. S. et al. Resolving the 3D Landscape of Transcription-Linked Mammalian Chromatin Folding. Mol. Cell 78, 539–553.e8 (2020).

37. Guo, Y. et al. Chromatin jets define the properties of cohesin-driven in vivo loop extrusion. Mol. Cell 82, 3769–3780.e5 (2022).

38. Hosoya, T. et al. Global dynamics of stage-specific transcription factor binding during thymocyte development. Sci. Rep. 8, 5605 (2018).

39. Ikawa, T. et al. Conversion of T cells to B cells by inactivation of polycomb-mediated epigenetic suppression of the B-lineage program. Genes Dev. 30, 2475–2485 (2016).

40. Ramani, V., Qiu, R. & Shendure, J. High Sensitivity Profiling of Chromatin Structure by MNase-SSP. Cell Rep. 26, 2465–2476.e4 (2019).

41. Sept, C. E. et al. High-resolution CTCF footprinting reveals impact of chromatin state on cohesin extrusion. Nat. Commun. 16, 4506 (2025).

42. Farley, E. K. et al. Suboptimization of developmental enhancers. Science 350, 325–328 (2015).

43. Crocker, J. et al. Low affinity binding site clusters confer hox specificity and regulatory robustness. Cell 160, 191–203 (2015).

44. Bruno, L. et al. Selective deployment of transcription factor paralogs with submaximal strength facilitates gene regulation in the immune system. Nat. Immunol. 20, 1372–1380 (2019).

45. Lim, F. et al. Affinity-optimizing enhancer variants disrupt development. Nature 626, 151–159 (2024).

46. Ou, F. et al. Optimization of the Irf8 +32-kb enhancer disrupts dendritic cell lineage segregation. Nat. Immunol. 25, 2043–2056 (2024).

47. Picelli, S. et al. Full-length RNA-seq from single cells using Smart-seq2. Nat. Protoc. 9, 171–181 (2014).

48. Abdennur, N. & Mirny, L. A. Cooler: scalable storage for Hi-C data and other genomically labeled arrays. Bioinformatics 36, 311–316 (2020).

49. Durand, N. C. et al. Juicer Provides a One-Click System for Analyzing Loop-Resolution Hi-C Experiments. Cell Syst. 3, 95–98 (2016).

50. Matthey-Doret, C. et al. Computer vision for pattern detection in chromosome contact maps. Nat. Commun. 11, 5795 (2020).

51. Lun, A. T. & Smyth, G. K. diffHic: a Bioconductor package to detect differential genomic interactions in Hi-C data. BMC Bioinformatics 16, 258 (2015).

52. Serizay, J. et al. Orchestrating chromosome conformation capture analysis with Bioconductor. Nat. Commun. 15, 1072 (2024).

53. Open2C et al. Cooltools: Enabling high-resolution Hi-C analysis in Python. PLoS Comput. Biol. 20, e1012067 (2024).

54. Chen, S., Zhou, Y., Chen, Y. & Gu, J. fastp: an ultra-fast all-in-one FASTQ preprocessor. Bioinformatics 34, i884–i890 (2018).

55. Langmead, B. & Salzberg, S. L. Fast gapped-read alignment with Bowtie 2. Nat. Methods 9, 357–359 (2012).

56. Quinlan, A. R. & Hall, I. M. BEDTools: a flexible suite of utilities for comparing genomic features. Bioinformatics 26, 841–842 (2012).

57. Xu, W. et al. CoolBox: a flexible toolkit for visual analysis of genomics data. BMC Bioinformatics 22, 489 (2021).

58. Robinson, J. T. et al. Integrative genomics viewer. Nat. Biotechnol. 29, 24–26 (2011).

